# Transcription-associated topoisomerase activities control DNA-breaks production by G-quadruplex ligands

**DOI:** 10.1101/2020.02.18.953851

**Authors:** A. Pipier, M. Bossaert, J.F. Riou, C. Noirot, L-T. Nguyễn, R-F. Serre, O. Bouchez, E. Defrancq, P. Calsou, S. Britton, D. Gomez

**Affiliations:** Institut de Pharmacologie et Biologie Structurale, IPBS, Université de Toulouse, CNRS, UPS, Toulouse, France; Equipe Labellisée Ligue Contre le Cancer 2018, Toulouse, France; Structure et Instabilité des Génomes, Muséum National d’Histoire Naturelle, CNRS, INSERM, Paris, France; INRAE, US 1426, GeT-PlaGe, Genotoul, Castanet-Tolosan, France; Département de Chimie Moléculaire, UMR CNRS 5250, Université Grenoble Alpes, Grenoble 38058, France

**Keywords:** G-quadruplex, CX5461, Topoisomerase 2, DNA-breaks, Transcription

## Abstract

G-quadruplexes (G4), non-canonical DNA structures, are involved in several essential processes. Stabilization of G4 structures by small compounds (G4 ligands) affects almost all DNA transactions, including telomere maintenance and genomic stability. Here, thanks to a powerful and unbiased genetic approach, we identify topoisomerase 2-alpha (TOP2A) as the main effector of cell cytotoxicity induced by CX5461, a G4 ligand currently undergoing phase I/II clinical trials. This approach also allowed to identify new point mutations affecting TOP2A activity without compromising cell viability. Moreover, based on cross-resistance studies and siRNA-based protein depletion we report that TOP2A plays a major role in cell cytotoxicity induced by two unrelated clastogenic G4 ligands, CX5461 and pyridostatin (PDS). We also report that cytotoxic effects induced by both compounds are associated with topoisomerase 2-mediated DNA breaks production. Finally, we show that TOP2-mediated DNA breaks production is strongly associated with RNA Pol II-dependent transcription and is countered by topoisomerase 1 (TOP1). Altogether our results indicate that clastogenic G4 ligands act as DNA structure-driven TOP2-poisons at transcribed regions bearing G-quadruplex structures.

## Introduction

Over the last years, accumulating evidence indicates that transcription is a major source of genomic instability (1, 2). Transcription-dependent DNA double-stranded breaks (DSBs) are mainly associated with RNA-Polymerase (RNA-Pol) arrests provoked by different non-exclusive factors including DNA torsional stress, inhibition of transcription elongation and formation of secondary structures, such as G-quadruplexes (G4) and R-loops (3–5). G4 are four-stranded secondary structures formed at guanine-rich tracts (6). Present throughout the human genome (7–9), G4 have been associated with spontaneous DNA breaks, hotspots for chromosomal translocations and several human syndromes (10–12). In proliferating cells, G4 act as replication fork barriers, provoking fork collapses, the activation of the DNA damage response and the induction of replication-dependent DSBs (13). However, increasing evidence indicates also a significant impact of G4 structures on genomic stability through transcription-dependent processes (5, 14–16).

From yeast to humans, mapping of DSBs at base-pair resolution identifies G4 structures as a critical factor promoting spontaneous DSBs (17). G4 mapping in the human genome shows a significant enrichment of these structures within promoter and 5’ UTRs regions of highly transcribed genes, and several *in vitro* and cellular studies show that the stabilization of G4 structures by small compounds, G4 ligands, represses transcription of genes containing G-rich tracts (7, 8, 10). While during transcription G4 structures located on template DNA could act as physical barriers blocking RNA-Pol II progression, the formation of G4 on the opposite strand could promote and stabilize secondary structures that block transcription elongation (3). For instance, genome-wide analyses of G4 motifs in human cells indicate that these structures are highly correlated with RNA-Pol II pausing sites and R-loop-forming regions, two different factors promoting RNA-Pol II arrests and transcription-dependent DNA breaks (17–19). Altogether, while numerous studies suggest a preponderant role of G4 in the formation of spontaneous transcription-dependent DNA breaks, it is not known how are DNA breaks formed and what are the cellular factors involved in this process. In eukaryotic cells, among the different DNA topoisomerases identified, two enzymes are required to resolve topological stresses (for a review see (20)). These enzymes relax topological constraints through the formation of transient single stranded (DNA topoisomerase I, TOP1) or double stranded DNA breaks (DNA topoisomerases II, TOP2), in which the enzymes are covalently linked to the DNA backbone. During transcription TOP1 protein removes both negative and positive supercoils induced by RNA Pol progression. TOP1 plays a critical role in suppressing genome instability mediated by non-canonical secondary structures such as G4 and R-loops that are promoted by transcription (21–25). In humans, TOP2 activity is supported by two isoenzymes TOP2 alpha (TOP2A) and TOP2 beta (TOP2B), that are encoded by two different genes. TOP2A plays key roles in DNA replication and chromosome segregation while TOP2B is mainly associated with transcription (20, 26, 27). TOP2 are poisoned by small molecules that trap the transient topoisomerase 2-DNA complex, known also as “covalent complex” (TOP2cc) during the enzyme catalytic cycle (28–30). The repair of TOP2cc requires a sequential process consisting of the removal of TOP2 protein from DNA through proteolytic (31–33) or nucleolytic degradation (34, 35) and the repair of the resulting DSB by Non-Homologous End Joining (NHEJ) (36) or Homologous Recombination (HR), respectively (37).

In this study, we show that the cell cytotoxicity induced by two potent and selective G4 ligands, CX5461 and pyridostatin (PDS) correlates with the formation of transcription dependent DNA breaks that are produced through a mechanism mediated by TOP2 and countered by TOP1 through the removal of RNA Pol II-dependent DNA topological stress and the release of RNA Pol II complex from transcription pausing sites.

## Results

### TOP2A mutations confer resistance to clastogenic G4 ligands CX5461 and pyridostatin

Currently in Phase I/II clinical trials for cancer treatments (38), the chemical compound CX5461, was firstly described as an RNA Pol I inhibitor (39). Although cytotoxic effects induced by this compound have been related to rDNA-transcription inhibition (40), CX5461 was also shown to be a potent G4 stabilizer and to provoke rapid induction of DSBs (41). In order to better define how CX5461 mediates its cytotoxicity we adopted an unbiased approach based on the selection and characterization of cells resistant to this drug. To do this, human near haploid HAP1 cells were randomly mutagenized with Ethyl Methane Sulfonate (EMS) based on previous work (42) and clones resistant to a lethal CX5461 concentration of 0.3 µM were isolated (CXR clones). Resistance of seven of these clones to CX5461 was further confirmed by cell survival assays showing IC_50_ values for CXR clones ranging from 0.223 to 0.335 µM of CX5461, corresponding to an average eight-fold increase in the IC_50_ value compared to the 0.033 µM IC_50_ of wild-type HAP1 cells (WT, see Supplementary table 1 and Figure 1a). In addition, CXR clones do not show cross-resistance to the efflux-pumps substrate nocodazole (43) indicating that multi-drug resistance (MDR) is not responsible for the observed resistance to CX5461 (Supplementary Figure 1a).

**Figure 1:**
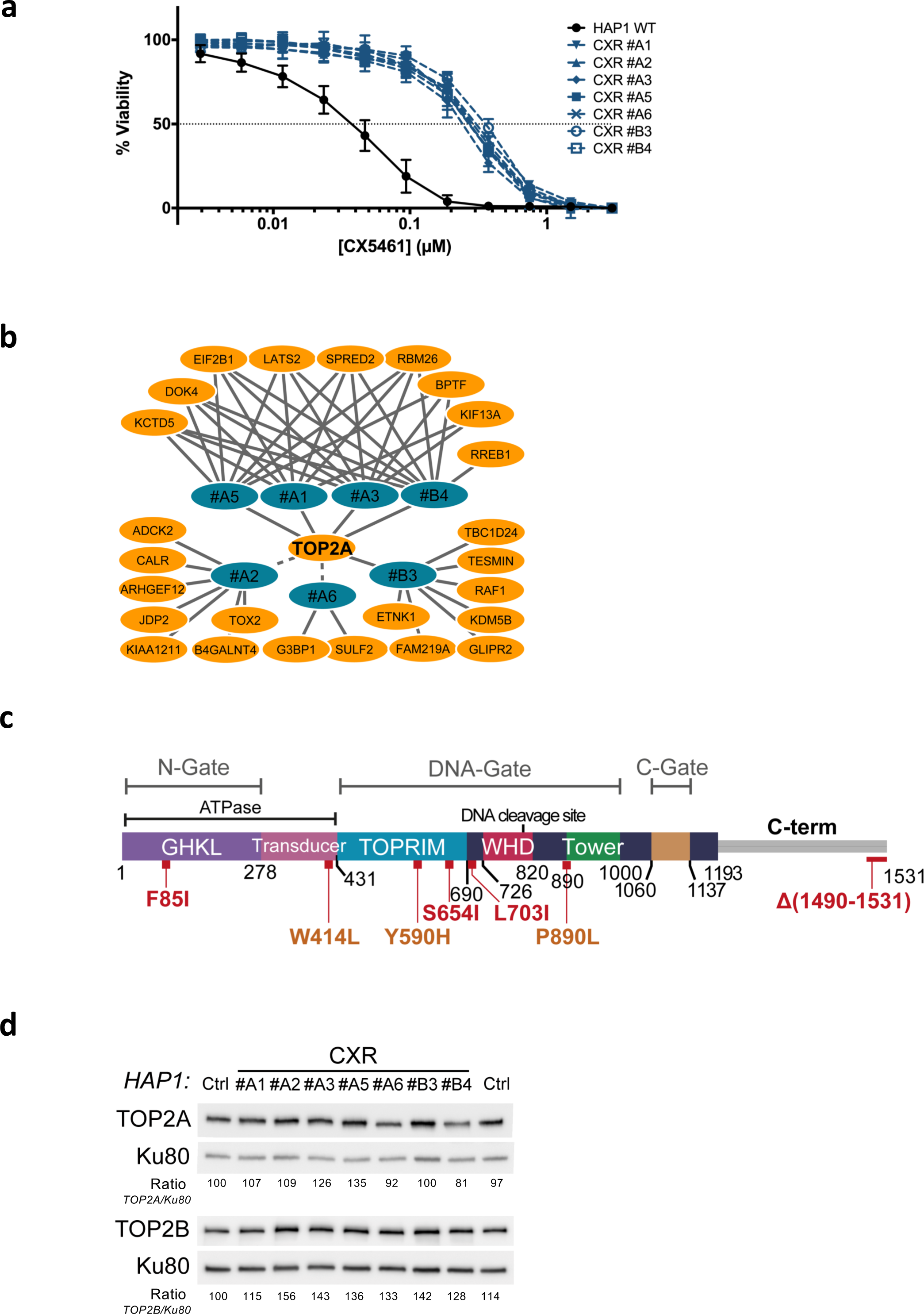
Role for TOP2A in the cell toxicity of CX5461. **a**, Viability assay on WT and seven CX5461 resistant (CXR) HAP1 clones treated with CX5461. Error bars represent SD from the means, *n* ≥ 3 independent experiments. **b**, Representation of genes with non- and mis-sense mutations identified in CXR clones. Solid and dashed lines represent respectively mutations characterized through an unbiased or manual analysis of RNA-seq data. **c**, Linear schematic of TOP2A domains. Each domain is labeled and described by bordering residue numbers. TOP2A mutations present in CX5461 or F14512 resistant clones are indicated in red or orange respectively. **d**, Immunoblotting analysis of whole-cell extracts from WT (Ctrl) and CXR HAP1 cells. Relative protein levels of TOP2A and TOP2B were quantified, normalized to KU80 level, and set to 100 in Ctrl cells.

Inspired by previous work (44), we analysed the selected CXR clones through a global RNA-sequencing approach (RNA-seq) to identify non- and mis-sense mutations in coding genes that could account for the observed resistance. Around 8 genes per clones were found with non- or mis-sense mutations through this approach (Figure 1b and Supplementary table 2). Unbiased analysis of genes found mutated in several clones revealed that each clone, except CXR#A2 and CXR#A6, carried a homozygous mutation in the *TOP2A* gene, encoding for the TOP2A protein (Figure 1b). Manual analysis of the sequencing data for the *TOP2A* gene confirmed these mutations and revealed that the CXR#A2 clone had the S654I mutation, while the CXR#A6 clone carried a homozygous mutation of the first nucleotide of the last intron, resulting in intron retention and replacement of the 42 last TOP2A amino acids, carrying its nuclear localisation signal (NLS), by 18 unrelated amino acids (Supplementary Figure 1b). From these analyses, *TOP2A* emerged as the only gene with coding mutations in all resistant clones. Four clones had a mutation in the ATPase domain (F85I), while two had mutations in the DNA binding region (S654I and L703I) (Figure 1c). None of these mutations were previously reported. Immunoblotting analysis revealed that none of the identified TOP2A resulted in loss of TOP2A protein (Fig 1d), in agreement with its essential function in proliferating cells (45, 46). In addition, TOP2B expression level was unaffected in these clones. To test whether the catalytic activity of TOP2A protein was altered in CXR clones, we determined the sensitivity of WT and resistant HAP1 clones to the TOP2 poison etoposide, a chemotherapeutic drug that acts by stabilizing TOP2cc and the cytotoxicity of which is therefore dependent on TOP2 activity (47). All CXR presented a strong resistance to ETP with resistance indexes ranging from eight to nineteen-fold relative to control cells (Figure 2a and Sup Table 1). These results support that TOP2A point mutations of CXR clones reduce TOP2A activity. To confirm that, we adapted a heparin-based extraction protocol (48) to monitor by immunoblotting the accumulation of TOP2Acc following ETP treatments. In this assay, TOP2 not covalently attached to DNA is extracted by heparin in a soluble fraction, while TOP2cc is resistant to this procedure and can be analysed by immunoblotting on pellet fraction. As show in Supplementary Figure 2, point mutations present on clones CXR#A6 and CXR#A1 decreased the number of TOP2Acc.

**Figure 2:**
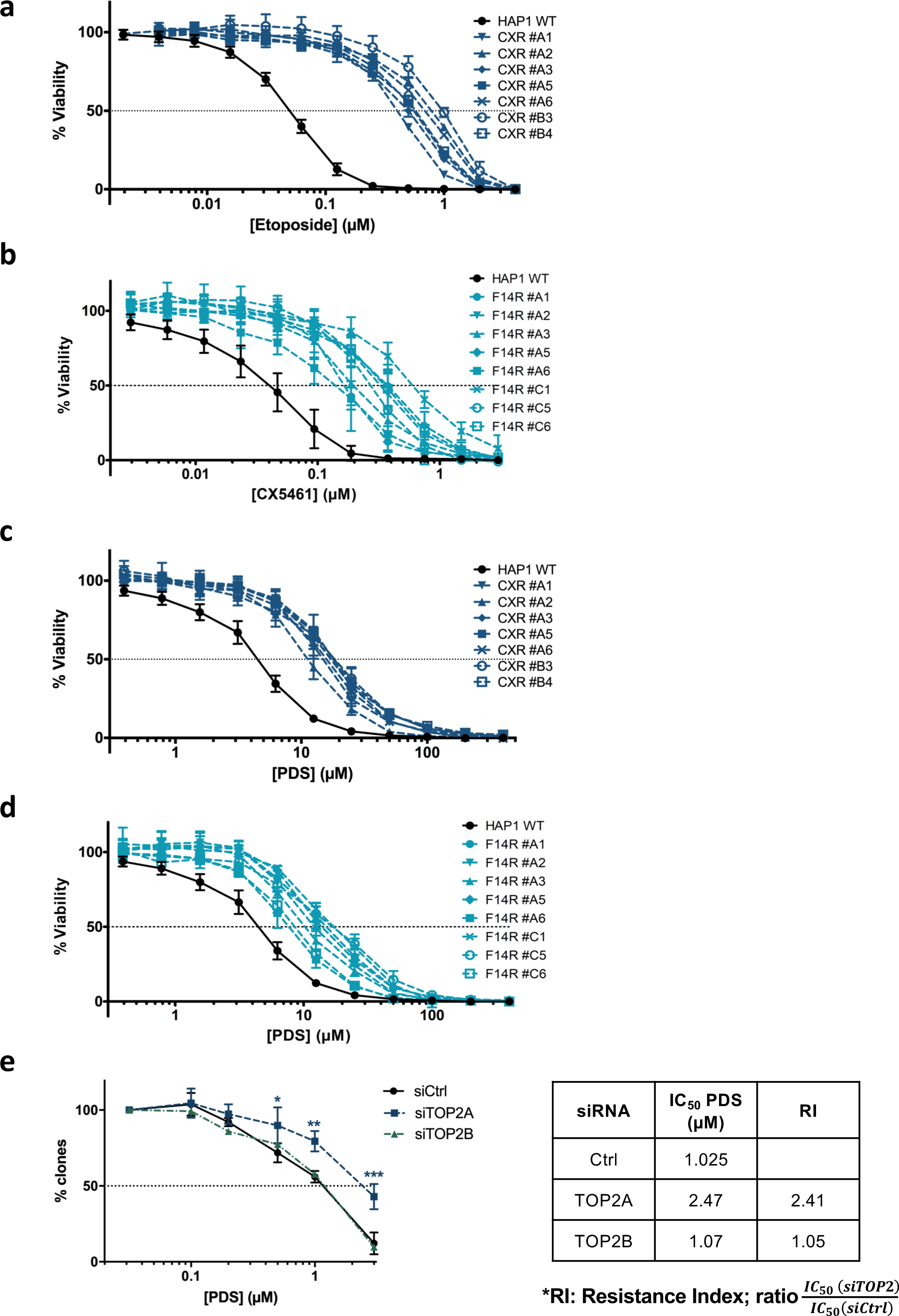
Impact of TOP2A mutations in the cell toxicity of topoisomerase poisons and G4 ligands. **a and c**, Viability assay and cross-resistance of CX5461 resistant cells (CXR) to the topoisomerase 2-poison etoposide and the G4 ligand PDS. **b and d**, Viability assay and cross-resistance of F14512 resistant cells (F14R) to the G4 ligands CX5461 and PDS. **e**, Graph representing cell survival as assessed by clonogenic assays on HeLa cells transfected with control (Ctrl), TOP2A or TOP2B siRNAs and treated with PDS. IC_50_ values of PDS on siRNA transfected cells are indicated in the right panel. The PDS resistance index (RI) was calculated as the ratio between IC_50_ values obtained for siRNA-TOP2 transfected cells and the IC_50_ value obtained for siRNA-Ctrl transfected cells. Error bars represent SD from the means, *n* ≥ 3 independent experiments. *P* values were calculated using an unpaired multiple Student’s *t* test. *: p<0.05; **: p<0.01; ***: p<0.001

In parallel, using the same genetic approach, we isolated F14R HAP1clones resistant to a lethal concentration of 30 nM F14512, a potent and selective TOP2A poison (49). Targeted sequencing of TOP2A cDNA in seven F14R clones revealed TOP2A mutations for five of them, confirming that TOP2A is the main mediator of F14512 cytotoxicity. One mutation was found in the transducer domain (W414L), while the two others lied in the DNA binding domain (Y590H and P890L, Figure 1c). These three mutations are different from the ones found in CXR clones, which could support that despite both acting though TOP2A, CX54161 and F14512 have different way to interfere with TOP2A activity. Strikingly, cell survival assays clearly demonstrate that all of F14R cells show cross-resistance to the CX5461 molecule (Figure 2b), with IC_50_ values for CX5461 3.6 to 16.8-fold higher than the IC_50_ value of CX5461 for WT cells (Figure 2b and Sup table 1) supporting that the TOP2A mutations found in F14R clones also confer resistance to the G-quadruplex ligand CX5461.

To investigate whether CX-resistance induced by TOP2A mutations was related to G4 stabilization, we tested CXR mutants for their cross-resistance to pyridostatin (PDS) one of the most potent and selective G4-ligands described so far (50). In cells, PDS treatment, similarly to CX5461, induces a rapid accumulation of DSBs, but in contrast to the CX5461, PDS has not been reported to affect RNA pol I activity (14). Cell survival assays showed that all CXR clones present a high resistance to PDS treatment with IC_50_ values for PDS 2.7 to 4.4-fold higher than the IC_50_ value of PDS for WT cells (Figure 2c and Sup table 1). More remarkably, cross-resistance studies established that all of F14R clones are also resistant to PDS (Figure 2d). Finally, to confirm and generalize the role of TOP2A protein in cellular cytotoxicity induced by G4 ligands in human cells, we monitored through clonogenic assays the survival to PDS of HeLa cells transfected with control siRNA or siRNA against TOP2A or TOP2B. As shown in Figure 2e, TOP2A knockdown results in a significant resistance to PDS as compared to control conditions while TOP2B depletion does not affect PDS cytotoxicity. Altogether, these results indicate that the catalytic activity of the TOP2A protein plays a major role in the cytotoxic effect induced by CX5461 and PDS compounds, two potent G4 stabilizers.

### Rapid accumulation of DNA double-strand breaks upon G4-ligands treatment depends on TOP2 proteins

In human cells, short treatments with PDS or CX5461 induce rapid production of DSB markers γH2AX and 53BP1 foci (14, 41), Figure 3a and Supplementary Figure 3a). G4-dependent γH2AX foci production is strongly increased in cells incubated with the DNA-PKcs inhibitor NU7441 (DNA-PKi, Supplementary Figure 3b), indicating that a substantial number of G4-induced DNA breaks are repaired through the NHEJ pathway, the major DSB repair pathway in human cells (51). Considering the main contribution of DSBs to the cytotoxic effect of several anticancer agents and having shown that TOP2A activity determines the cytotoxic effect of G4 ligands (Figures 1&2), we evaluated TOP2A role in DSB production upon PDS and CX5461 treatments. First, in CXR cells carrying different mutations on TOP2A protein (clones CXR #A1, #A2 and #A6), γH2AX production was significantly reduced as compared to control cells (Figure 3a). In addition, γH2AX production in HeLa cells transfected by siRNA against TOP2A was abolished upon PDS treatment and strongly reduced upon CX5461 treatment, as compared to cells transfected with control siRNA (Figure 3b and Supplementary Figure 3c). In contrast, TOP2B depletion did not impact γH2AX production upon PDS treatment while it reduced γH2AX production upon CX5461 to a similar extend than TOP2A depletion. This result indicated that, while TOP2A is the main contributor to DSB induction on response to PDS, both TOP2A and TOP2B are involved for CX5461. In agreement, simultaneous siRNA-mediated knock-down of TOP2A and TOP2B prevented γH2AX production by CX5461 (Figure 3c). Finally, to validate the role of TOP2 activities on DSBs production induced by PDS treatments, we studied the impact of the TOP2 catalytic inhibitor BNS-22 (52) on DSB production by PDS. Pre-incubation with BNS-22 significantly decreased the number of γH2AX foci in HeLa cells treated with PDS, thereby confirming the major role of TOP2 activities in the production of DNA breaks following G4 ligand treatments (Figure 3d).

**Figure 3:**
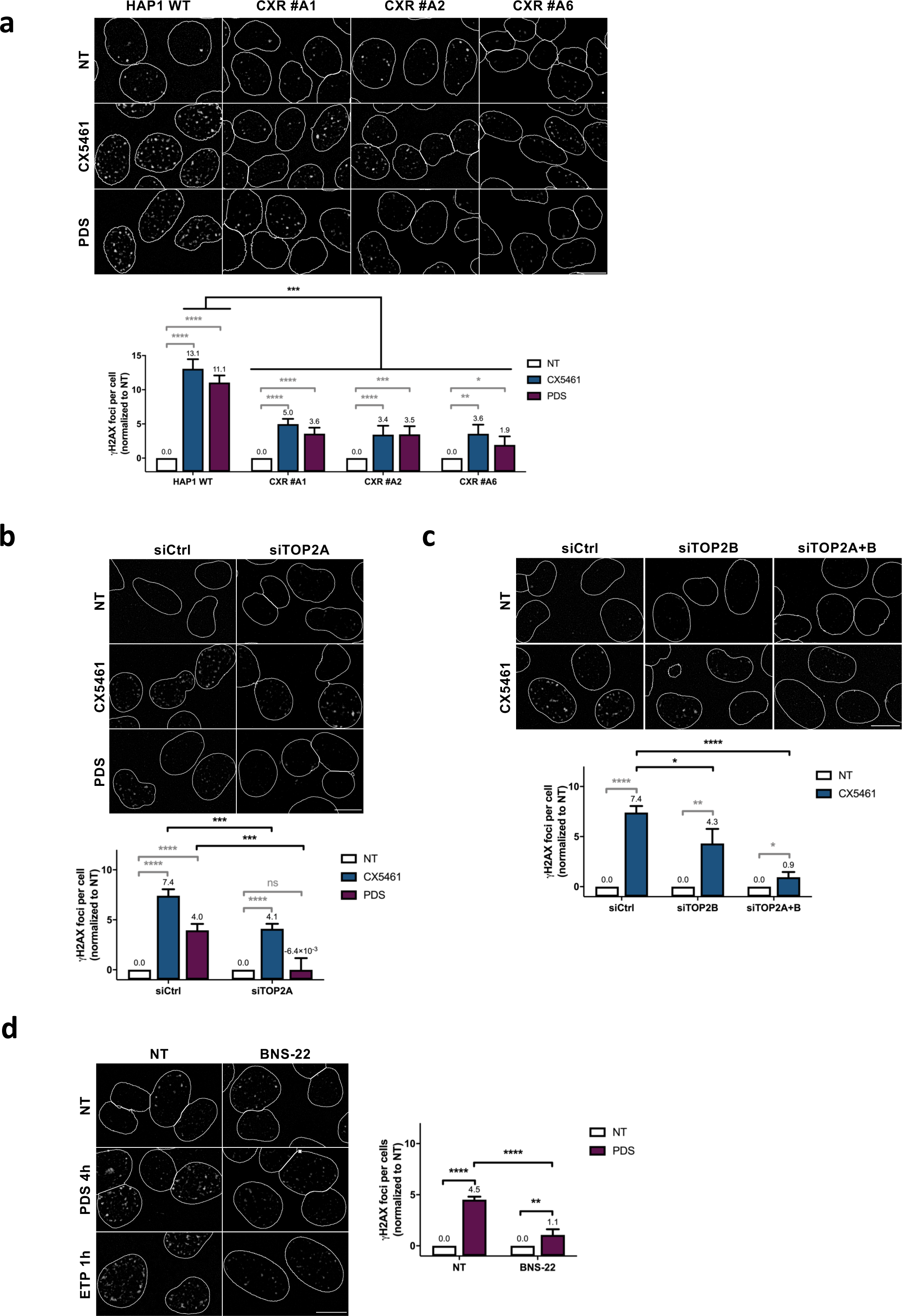
Role of topoisomerase 2 proteins in DNA breaks production by G4 ligands CX5461 and PDS. Quantification and representative images of γH2AX foci fluorescence signal (grey) detected in: HAP1 WT and CXR cells (**a**) or in HeLa cells transfected with control (Ctrl), TOP2A and/or TOP2B siRNAs and treated with PDS or CX5461 (**b and c**). **d**, Quantification and representative images of γH2AX foci detected after PDS or Etoposide treatment in HeLa cells pre-treated with the topoisomerase 2 catalytic inhibitor BNS-22. For all the experiments, cells were incubated with CX5461 (0.2 µM) and PDS (20 µM) for 4 hours. For BNS-22 experiment the inhibitor (5 µM) was added 30 min prior to addition of PDS. Quantification of γH2AX foci per cell was performed as described in Methods on *n* > 165, *n*>105, *n*>105 and *n*>101 nuclei for each condition respectively in **a, b, c** and **d**. Error bars represent SD from the means, *n* ≥ 3 independent experiments. *P* values were calculated using an unpaired multiple Student’s *t* test. ns: p>0.05; *: p<0.05; **: p<0.01; ***: p<0.001; ****: p<0.0001

### TOP2-dependent DSBs induced by G4 ligands are transcription-dependent

In human cells, the appearance of DNA damage signals after PDS treatment was shown to be independent on cell cycle status as PDS-induced DSB markers are observed in G1, G2 and S cell cycle phases (Supplementary Figure 4a and (14)). Moreover, simultaneous incubation with PDS and the DNA-base analogue EdU (5-ethynyl-2′-deoxyuridine), used to visualize DNA synthesis, shows that the induction of DNA damage by PDS can occurs independently of the DNA replication process ((14) and Supplementary Figure 4b). Furthermore, the average of PDS-induced DSB markers in EdU negative cells compared to the average of these markers in the total cell population indicates that DNA replication-independent processes are very efficient for the production of DSBs by PDS (Supplementary Figure 4b). Indeed, in HeLa cells, production of DNA damage by PDS was almost completely abolished by inhibition of RNA Pol II-dependent transcription by DRB (5,6-dichloro-1-b-D-ribofuranosylbenzimidazole), an inhibitor of critical phosphorylations of RNA Pol II C-terminal domain (53). DRB also reduced DSB production upon CX5461 treatment, albeit to a lesser extent than with PDS (Figure 4a). In addition, and in line with this finding, immunofluorescence studies with the G4-specific antibody BG4 show that the inhibition of RNA Pol II-dependent transcription does not impede the accumulation of G4 structures caused by PDS-treatments (Supplementary Figure 4c). These data argue for a key role RNA-Pol II transcription elongation in the production of DSBs following the stabilization of G4 structures by G4 ligands.

**Figure 4:**
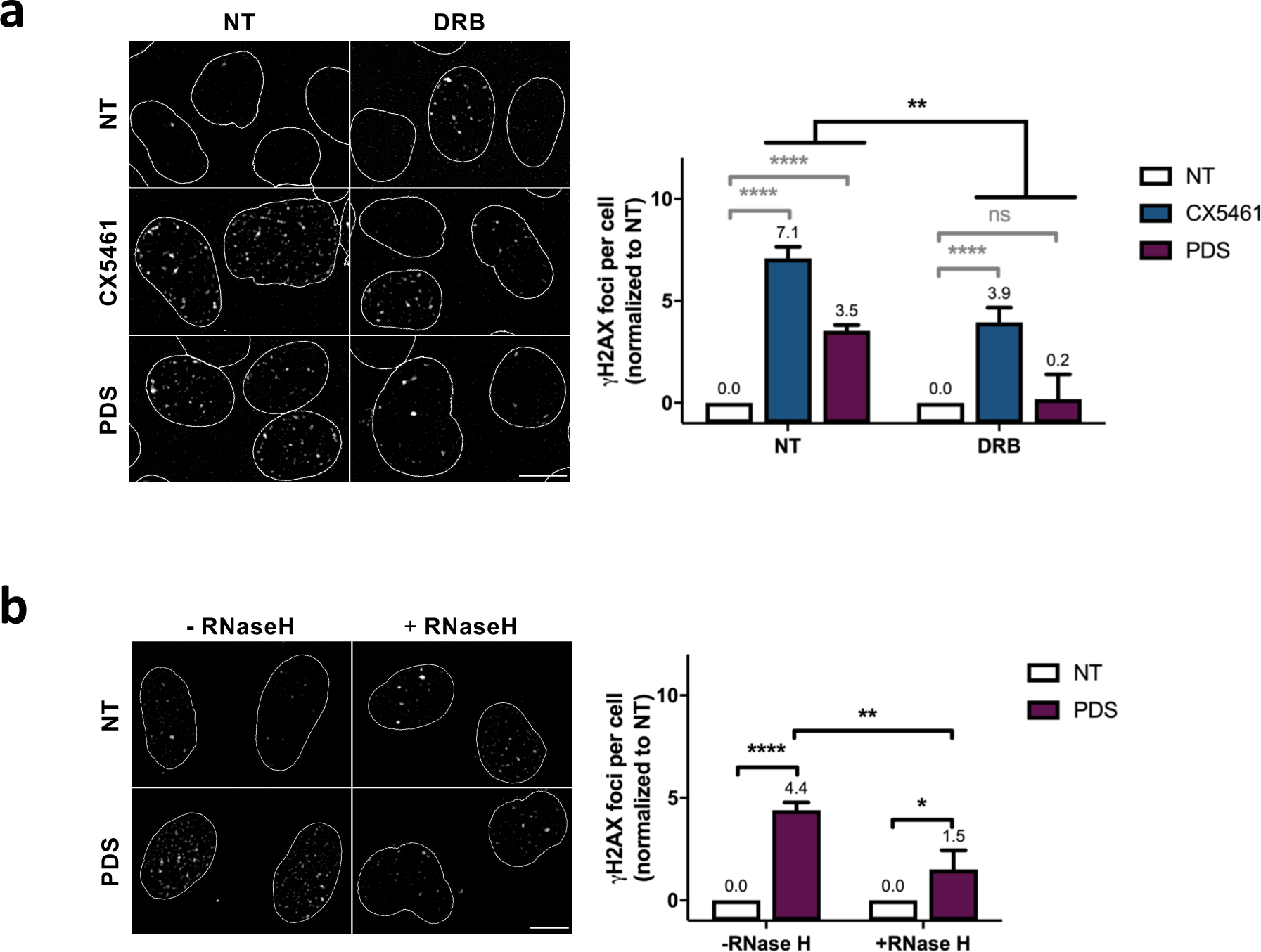
Role of RNA Pol II-dependent transcription in DNA breaks production by G4 ligands CX5461 and PDS. Quantification and representative images of γH2AX foci fluorescence signal (grey) detected in: HeLa cells pre-treated with the RNA Pol II inhibitor DRB prior addition of CX5461 (0.2 µM) and PDS (20 µM) for 4 hours **(a)** and in RNaseH1-mCherry U2OS expressing cells for 8 hours **(b).** DRB (100 µM) was added 1 hour before CX5461 or PDS addition. RNaseH1-mCherry expression in U2OS cells was induced with 2.5 µg/mL doxycyclin 14 hours prior to PDS treatment. γH2AX foci per cell was performed as described in Methods on *n* > 165 nuclei for each condition in **a** and *n* > 42 nuclei for each condition in **b**. Error bars represent SD from the means, *n* ≥ 3 independent experiments. *P* values were calculated using anunpaired multiple Student’s *t* test. ns: p>0.05; *: p<0.05; **: p<0.01; ***: p<0.001; ****: p<0.0001

R-loops have been associated with the production of transcription-dependent DSBs and genomic instability. Thus, we investigated the role of these structures on the DNA damage production induced by CX5461 and PDS compounds by using U2OS cell line expressing the E. coli RNaseHI under the control of a doxycycline inducible promoter (54). Upon RNAseHI expression, we observed a net decrease of DNA damage signals induced by PDS (Figure 4b), indicating that transcription-associated R-loops structures contribute to the formation of DNA breaks following G4 stabilization Altogether, these results indicate that RNA-Pol II transcription elongation plays a key role in the production of DNA damage by both G4 ligands.

### Top2-dependent DSBs induced by G4 stabilizers are countered by Top1

In cells, topological stresses provoked by transcriptional elongation are principally relieved by TOP1 (55, 56). These topological changes provoke both DNA melting (negative supercoiling) behind the RNA pol II complex facilitating the formation of non-B DNA structures, such as G4 (21, 25, 57) and the accumulation of positive supercoiling in front of the RNA complex that act as barriers to transcriptional elongation (58, 59). Since we observed that DSBs induced by G4 ligands are mainly dependent on transcription, we evaluated the role of TOP1 enzyme in the cellular response to these compounds. Strikingly, RNA-silencing mediated depletion of TOP1 protein in human cells caused a significant increase in DNA damage induced by CX5461 and PDS (Figure 5a) that are completely dependent of transcription as DRB pre-treatments completely abrogated PDS-induced DNA damage signals in TOP1 depleted cells (Figure 5b). Moreover, EdU staining indicated that enhanced DNA damage production in TOP1 knockdown cells is not restricted to the S phase of the cell cycle (Supplementary Figure 5). In agreement with these data, the depletion of TOP1 in HeLa cells caused a significant increase in the cytotoxic effect of PDS (3-fold decrease in the IC_50_) that can be reverted by a DRB pre-treatment (Figure 5c). Altogether, these results strongly suggest that the accumulation of topological stresses provoked by RNA Pol II-dependent DNA transcription in the absence of the TOP1 protein are key factors in the mechanism of DNA damage production and cell toxicity promoted by G4 ligands. Consistent with these findings, immunofluorescence studies show that TOP1 knockdown provokes a significant increase of BG4 signals in human cells (Figure 5d) that could result from the accumulation of transcription-dependent negative supercoiling caused by TOP1 depletion. In addition, since a cumulative effect of PDS and TOP1 depletion on BG4 signals is not observed, we conclude that TOP1 depletion has a major impact on the processes leading to G4s formation. Furthermore, kinetics studies of γH2AX production in PDS-treated cells show that TOP1 depletion significantly accelerates the formation of PDS-induced DNA damage signals compared to control cells (Figure 5e).

**Figure 5:**
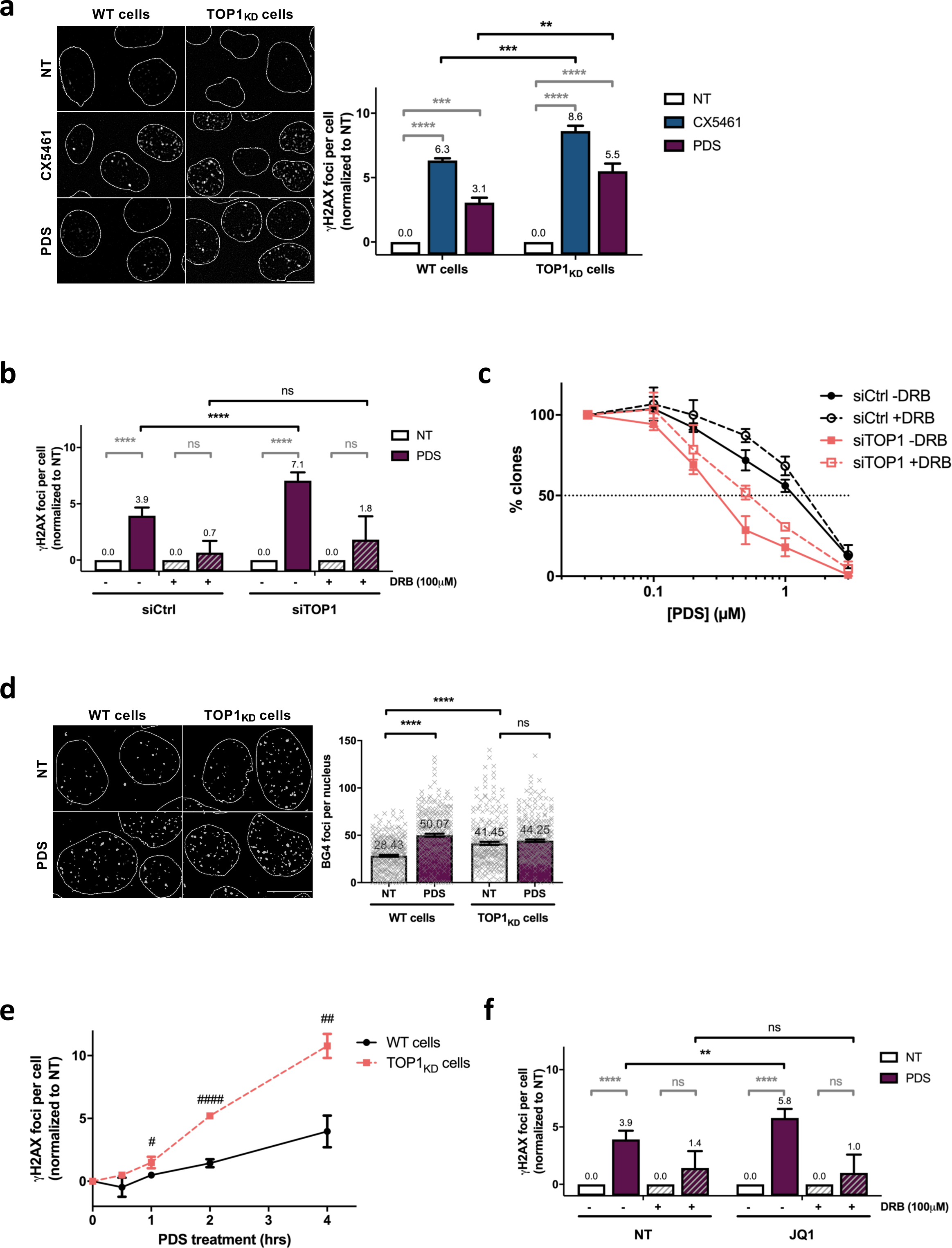
TOP1 protein counteract TOP2-dependent DSBs induced by G4 stabilizers. **a**, Quantification and representative images of γH2AX foci fluorescence signal (grey) detected in HeLa cells transfected with control (Ctrl), or TOP1 siRNAs and treated with PDS or CX5461. **b**, Quantification of γH2AX signals in HeLa cells transfected with control (Ctrl), or TOP1 siRNAs and pre-treated with the RNA Pol II inhibitor DRB. DRB (100 µM) was added 1 hour before PDS addition. **c**, Cell survival assay as assessed by clonogenic assay on HeLa cells transfected with control (Ctrl), or TOP1 siRNAs and treated with PDS and DRB. For clonogenic assays PDS and DRB treatment were performed as described in Methods. **d**, Quantification and representative images of ΒG4 foci fluorescence signal (grey) detected in HeLa cells transfected with control (Ctrl) or TOP1 siRNAs and treated with PDS (20 µM) for 4 hours. **e**, Kinetics studies of γH2AX foci formation in HeLa cells transfected with control (Ctrl), or TOP1 siRNAs following PDS (20 µM) treatments. **f**, Quantification of γH2AX signals in HeLa cells treated with PDS and the BRD4 inhibitor JQ1. JQ1 (2 µM) was added 1h prior to addition of PDS. Quantification of γH2AX foci per cell was performed as described in Methods on *n* > 110, *n* > 181, *n* > 174 and *n* > 117 nuclei for each condition respectively in **a**, **b**, **e** and **f**. Error bars represent SD from the means, *n* ≥ 3 independent experiments. *P* values were calculated using a unpaired multiple Student’s *t* test. ns: p>0.05; *: p<0.05; **: p<0.01; ***: p<0.001; ****: p<0.0001. Quantification of BG4 foci per cell was performed as described in Methods on *n* > 71 nuclei for each condition. Error bars represent sem from the means, *n* = 3 independent experiments. *P* values were calculated using an unpaired Whelch’s *t* test. ns: p>0.05; *: p<0.05; **: p<0.01; ***: p<0.001; ****: p<0.0001.

In mammals, stimulation of TOP1 activity is required to release the RNA Pol II complex from pausing sites that are characterized by a prolonged and regulated polymerase arrest in a promoter proximal position. TOP1 activity at paused RNA Pol II is enhanced by the BRD4-dependent phosphorylation of RNA Pol II CTD (60). Transcriptional pausing has been associated with DSBs formation and TOP2 activities (61–64). Furthermore, G4 motifs correlate with promoter-proximal transcriptional pausing in human genes (19). Interestingly, we show through immunofluorescence analysis that the inhibition of BRD4 activity by JQ1, a potent inhibitor of the BET family of bromodomain, provokes a significant increase of DNA breaks induced by PDS (Figure 5f), indicating that RNA pause release and TOP1 stimulation play an important role in the response to PDS. Altogether, our results strongly suggest that TOP1 antagonizes G4 ligands action through a transcription-dependent mechanism that is related to RNA polymerase pausing.

## Discussion

In this study, thanks to an unbiased genetic approach, we identified the TOP2A protein as the main effector of cell cytotoxicity induced by CX5461, a compound envisioned as an anticancer treatment and currently undergoing phase I/II clinical trials (38). Molecular characterization of resistant cell lines showed that single point mutations affecting TOP2A activity, as demonstrated both by cross resistance with etoposide and molecular approaches based on heparin extraction, significantly protect human cells from CX5461 induced cytotoxicity.

Our study highlights the strength of the genetic approach we applied here, relying on chemical mutagenesis in a haploid background as previously described (42). Indeed, despite TOP2A being essential in proliferating cells (45, 46), we were able to readily isolate CX5461 resistant clones carrying TOP2A mutations allowing direct identification of its crucial role in DSB induction upon G4 stabilisation, which would not have been possible through loss of function screens, for example using CRISPR/Cas9 or insertional mutagenesis. In addition, while this approach was aimed at identifying critical mediators of G4 cytotoxicity, it also provides novel mutations of TOP2A conferring resistance to both CX5461 and F14512 which could be useful to get insights into TOP2A biology. Thus, thanks to this genetic approach we were able to identified six-point mutations that are broadly distributed throughout the TOP2A coding sequence. Three mutations mapped in the DNA binding, one in the TOPRIM domain and two in the ATPase domain. A similar broad distribution of point mutations affecting TOP2A activity has been obtained through a complementation approach in yeast (65), indicating that, as observed for topoisomerase poisons, resistance to CX5461 can be obtained through different protein modifications and are not strictly dependent on the catalytic function. This is especially true for the mutation in the CXR#A6 clones which results in the expression of a TOP2A depleted from its terminal nuclear localisation sequence and therefore sequestered in the cytoplasm in interphase cells (Supplementary Figure 1b) while it can access DNA in mitosis.

Such a mutant would have been difficult to devise and to express at proper levels in complementation experiments (TOP2A overexpression is toxic, (66)). Two of the six amino acid changes in TOP2A identified in this work have been previously described and confer resistance to Vosaroxin a quinolone derivative that acts as a DNA intercalator and a topoisomerase II inhibitor (65) confirming that selected TOP2A point mutations through this genetic approach alter TOP2A activity.

A major finding of our work is that TOP2A protein mutations present in CXR cells also confer resistance to PDS, another potent and selective G4 ligand non-chemically related to CX5461 (50). The impact of TOP2A on cell cytotoxicity induced by PDS was further supported by survival assays that demonstrated that TOP2A siRNA mediated depletion drives PDS resistance in HeLa cells. Although CX5461 compound was initially identified as an RNA Pol I inhibitor (39) and TOP2A protein has been involved in rDNA transcription (67), we assumed that cellular resistance to CX5461 and PDS in TOP2A deficient cells was not directly related to rDNA transcription inhibition but most probably to the induction of DNA breaks through G4 stabilization. Consistent with this hypothesis, BRCA2 deficient cells, impaired for homologous-recombination DNA repair pathway, are highly sensitive to CX5461 and PDS (5, 41, 68, 69), indicating a major role of DNA repair mechanisms in the cellular response to both compounds. In contrast, BRCA2 ^−^/^−^ cells do not show increased sensitivity to BMH21, a very potent RNA Pol I inhibitor (41).

In agreement with these results, another finding in our study is that DNA breaks formation caused by short term treatments with PDS and CX5461 are mostly dependent on TOP2 proteins. Thus, siRNA mediated depletion of both isoforms or the inhibition of their catalytic activity by BNS-22 (52) provoke an almost complete loss of DNA breaks production induced by CX5461 and PDS treatments in human cells. Detailed analysis of the impact of two TOP2 isoforms in DNA break production following G4 ligands treatments show some differential contributions.

Thus, TOP2A depletion has a major impact in response to PDS, while both TOP2A and TOP2B are involved in CX5461-induced DNA breaks production in HeLa cells. Similar differences were also observed in response to transcriptional inhibition. Indeed, while DRB treatments, that specifically blocks RNA Pol II transcriptional elongation, completely abrogates DSBs formation induced by PDS, RNA Pol II inhibition provokes a significant but only partial reduction of DNA breaks production induced by CX5461 in HeLa cells. Altogether our results clearly demonstrate a major role of topoisomerase 2 activities in DNA breaks production induced by two clastogenic G4 ligands, CX5461 and PDS and a major contribution of RNA Pol II-dependent DNA transcription during this process.

The major contribution of TOP2A protein in PDS-dependent induced DNA breaks appears slightly surprising as a major role on transcription has been credited to TOP2B isoform (20, 26, 27, 70). In human cells, TOP2A is highly expressed in proliferating cells, and further up-regulated during S and G2 phases where it plays an essential role during replication and chromosome segregation (20, 26, 45, 46). However, TOP2A has also been implicated in transcription (67, 71) and its activity is also required for a maximal production of transcription-dependent DNA breaks induced by etoposide, a potent poison of topoisomerase 2 (72). Moreover, genome wide analysis of TOP2A cleavage sites show a significant enrichment of TOP2A on highly transcribed loci (33). Interestingly, elevated transcription levels have been shown to promote G4 formation that are favoured by negative superhelicity caused by the progression of RNA Pol complexes through DNA (57). In cells, topological stresses induced by RNA Pol II progression are mainly resolved by the TOP1 protein (73). In this study, we evidenced a major role of TOP1 protein in countering DNA breaks formation by CX5461 and PDS. Thus, we observed a dramatic increase of DNA breaks signals induced by both compounds in TOP1 depleted cells. Consistent with our data, TOP1 depletion in yeast drives genomic instability at highly transcribed G4 forming sequences (21, 25). Strikingly, DNA break formation upon PDS treatment in TOP1 knockdown cells was completely suppressed following TOP2A depletion (Supplementary Figure 6). This last result clearly confirms the preponderant role of TOP2A in the formation of transcription dependent DNA breaks following PDS treatments.

Essential role of topoisomerase enzymes in DNA break production following CX5461 and PDS treatments strongly suggests a TOP2-poisoning based mechanism. In cells, topoisomerase poisons stabilize normally transient topoisomerase 2-DNA complexes, promoting the formation of DSBs, one of the most deleterious DNA lesions for human cells (29, 30). Cellular resistance to topoisomerase 2 poisons is associated with reduced topoisomerases expression or with mutations affecting topoisomerase 2 catalytic activities (74). Consistent with the hypothesis that G4 ligands can also act as TOP2 poisons, in the present study we showed that single point mutations affecting TOP2A activity as well as the inhibition of TOP2 catalytic activities in human cells (BNS-22 treatments) strongly reduce DNA damage production induced by CX5461 and PDS. Dual *in vitro* activity of G4 ligands have been already reported (75, 76). Similarly, some TOP2 inhibitory molecules have been shown to possess G4 stabilization capacities (77–80).

Why G4 ligands-induced breaks are toxic? In cells, inhibition of DNA-PKcs activity dramatically increases the number of DNA break signals in PDS treated cells, demonstrating that an important number of PDS-induced DNA breaks are repaired through the NHEJ pathway. Strikingly, additional DNA breaks signals revealed under NHEJ deficiency seem independent of TOP2A as a similar increase of DNA breaks was observed in TOP2A deficient cells compared to control cells (Supplementary Figure 7). This result suggests that some breaks do not rely on TOP2A and are efficiently repaired by NHEJ while TOP2-dependent DNA breaks induced by G4 ligands are refractory to repair by NHEJ pathway, more likely supporting the toxicity of these molecules

Very recently, Bruno et al. through data mining, biochemical and cellular assays provided evidence that CX5461 mediates its cytotoxic effect through interference with TOP2 activity and not through inhibition of RNA polymerase I-dependent transcription (81). However, the underlying mechanism and the contribution of the TOP2A and TOP2B remained unclear. Here, we extent their observation to two chemically unrelated G4-ligands. Through identifying independent and original TOP2A mutations that compromise the catalytic activity of the protein, we definitively demonstrate that toxicity of these two G4-ligands require topoisomerase II activity that is responsible for DNA breaking at G-rich transcribed loci. In addition, we show that TOP1 counteracts G4-ligands clastogenic and toxic properties.

On the basis of the results obtained in this study, we propose a model in which G4 ligands CX5461 and PDS act as “G4-dependent TOP2 poisons” (Figure 6). In this model the interaction of both compounds with DNA is facilitated by DNA topological stress provoked by RNA Pol II-dependent transcription. G4 stabilization by G4 ligands in transcriptional active loci would provoke sustained RNA Pol II arrest mobilizing topoisomerase enzymes to resolve topological stresses and that at some loci may be poisoned at the vicinity of G4. Our model unifies the topoisomerase poisoning and G4-binding properties of these molecules in the new concept of DNA structure-driven topoisomerase poisoning at transcribed G-rich sequences.

**Figure 6:**
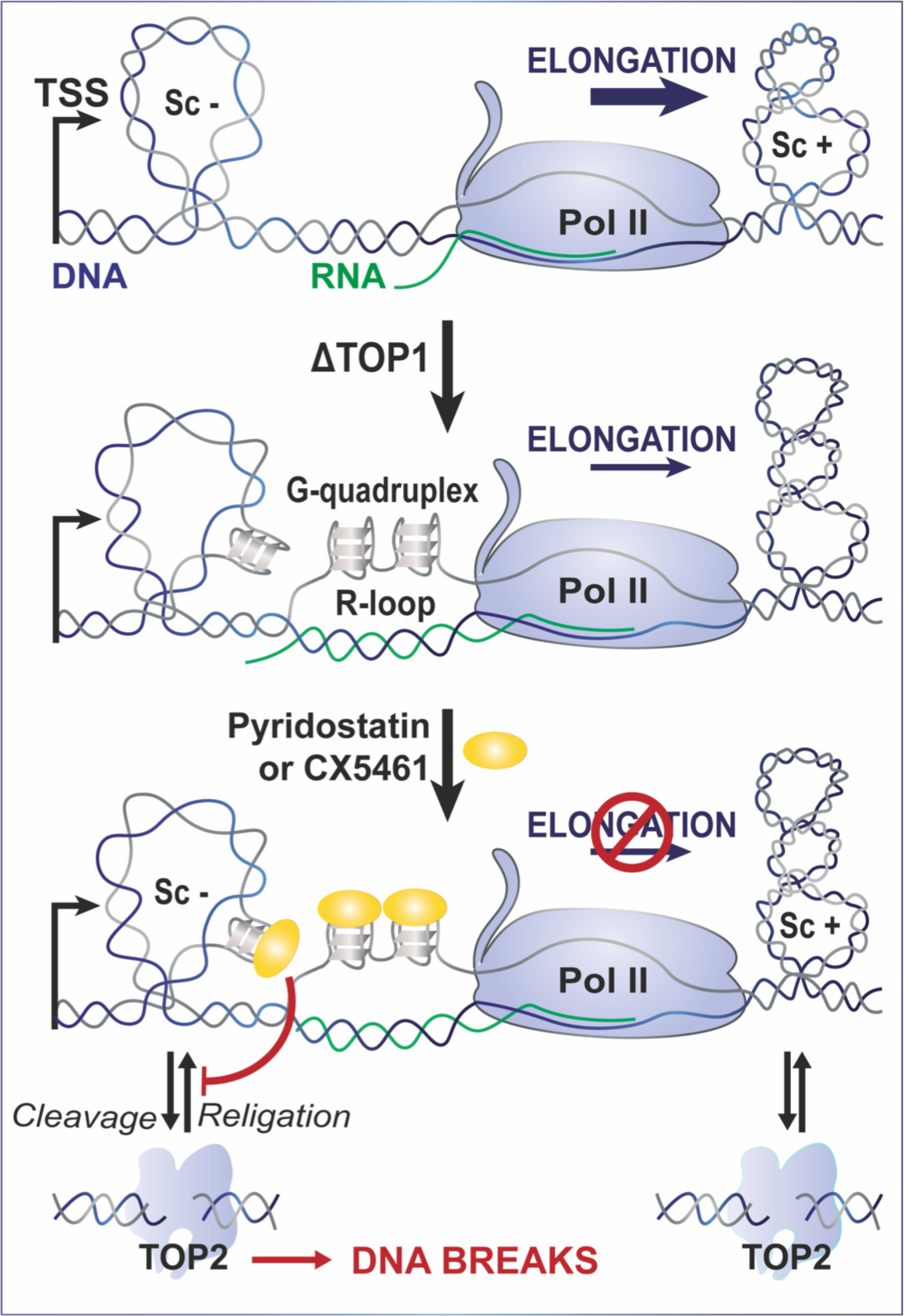
Proposed model for Topoisomerase 2-mediated DSBs on transcriptionally active loci containing G-quadruplex forming sequences. In this model the interaction of G4 ligands with DNA is facilitated by DNA topological stress provoked by RNA Pol II-dependent transcription and counteracted by TOP1 activity. G4 stabilization by G4 ligands in transcriptional active loci would provoke sustained RNA Pol II arrest mobilizing topoisomerase enzymes to resolve topological stresses and that at some loci may be poisoned at the vicinity of G4.

## Acknowledgments

This work was funded by grants from ANR (ANR-17-CE18-0002-01 and ANR-16-CE11-0006-01), Cancéropôle GSO Emergence funding “CX-Break” and La Ligue Nationale Contre le Cancer (Equipe Labellisée 2018). We are grateful to Emmanuelle Näser for technical assistance and the Imaging Core Facility TRI-IPBS, supported by ITMO Cancer (Alliance Nationale pour les Sciences de la Vie et de la Santé, National Alliance for Life Sciences and Health) within the Framework of the Cancer Plan. We thank “Région Midi-Pyrénées” for supporting the “Toulouse Réseaux Imagerie” platform. Patrick Calsou is a researcher from INSERM. This work was performed in collaboration with the GeT core facility, Toulouse, France (http://get.genotoul.fr), and was supported by France Génomique National infrastructure, funded as part of “Investissement d’avenir” program managed by Agence Nationale pour la Recherche (contract ANR-10-INBS-09).

## Material and Methods

### Cell culture conditions and treatments

All culture media were provided by Gibco and were supplemented with 10% fœtal bovine serum (Eurobio), 100U/mL penicillin (Gibco) and 100µg/mL streptomycin (Gibco). Cells were grown in humidified atmosphere with 5% CO2 at 37°C. HeLa and U2OS cells were grown in Dulbecco’s Modified Eagle Medium. HAP1 cells were cultured with Iscove’s Modified Dulbecco’s Medium. RPE1-hTERT cells were cultured with RPMI Media 1640 buffered with 0.3% Na(CO_3_)_2_. Expression of shTOPI in HeLa cells were induced with 5µg/mL doxycycline for 5 days before treatments and for 4 days before siTOP2A transfections (described below) and maintained during siRNA-mediated protein depletion. RNaseHI-mCherry expression in U2OS cells was induced with 2.5 µg/mL doxycyclin for 14 h.

For selection of HAP1 resistant clones, haploid HAP1 were isolated using cell sorting and 100.10^6^ haploid HAP1 were mutagenized by treatment with 300 µg/ml ethyl methane sulfonate (Sigma-Aldrich) for 3 days. After a one-week recovery, 0.5-1.10^6^ cells were seeded in 140 mm dishes. Plates were treated twice at a one-week interval with 0.3 µM CX5461 or 30 nM F14512 for 4 days. Around 10 days after the second treatment, individual clones (CXR and F14R) were isolated and used for further studies.

For immunofluorescence studies in U2OS, pyridostatin (PDS) was used at 20 µM for 8 hours. For immunofluorescence studies in HeLa and HAP1 cells, PDS and CX5461 were used respectively at 20 µM and 0.2 µM for 4 hours. 5-Ethynyl-2’-deoxyuridine (EdU) treatments were performed at 100 µM at the same time than PDS and CX5461 treatments and were maintained for the duration of experiments. For inhibitors, DNA-PKi (2 µM, NU7441) and JQ1 (2 µM, (+/-)-JQ1) were added to cells 1 h and BNS-22 (5 µM) was added to cells 30 min prior to pyridostatin and or CX5461 treatments. The transcription inhibitor DRB was used at 100 µM and added to cells 1 h prior to treatments. All inhibitors remained onto cells for the duration of the experiment.

| Chemical Compound | Furnisher | CAS number |
| --- | --- | --- |
| Pyridostatin | Sigma-Aldrich | 1085412-37-8 |
| CX5461 | Selleckchem | 1138549-36-6 |
| Doxycycline hydrochloride | Sigma-Aldrich | 10592-13-9 |
| DRB, 5,6- dichloro-1-b-D-ribofuranosylbenzimidazole | Sigma-Aldrich | 54-85-0 |
| DNA-PKi – NU7441 | Tocris | 503468-95-9 |
| Etoposide | Sigma-Aldrich | 33419-42-0 |
| BNS-22 | Sigma-Aldrich | 1151668-24-4 |
| Nocodazole | Sigma-Aldrich | 31430-18-9 |
| EdU, 5-Ethynyl-2'-deoxyuridine | Fisher-Scientific | nd |
| JQ1, (+/-)-JQ1 | Sigma-Aldrich | 1268524-69-1 |

### Plasmid and cell constructions

U2OS cells conditionally expressing the RNaseH1-mCherry fusion protein were previously described in (54). To construct HeLa cells conditionally expressing shRNAs against TOP1 mRNA, HeLa were infected with pLV-tTR-KRAB-Red and pLVTHM shTOP1 lentiviral particles. Individual clones from the transduced cell population were then isolated and selected for their capacity to downregulate TOP1 expression under treatment by the tetracyclin analog doxycyclin. pLV-tTR-KRAB-Red is a lentiviral vector encoding the transcriptional repressor tTR-KRAB fused to the DsRed fluorescent protein. pLVTHM is a lentiviral vector allowing conditional expression of an shRNA of interest under the control of the H1 promoter and the tetracyclin operator/repressor system (TetO/TetR). pLVTHM vector allowing conditional expression of an shRNA against TOP1 was obtained by inserting duplex oligonucleotides (5’-CGCGTCCCCGGACTCCATCAGATACTATTTCAAGAGAATAGTATCTGATGGAGTC CTTTTTGGAAAT-3’ and 5’-CGATTTCCAAAAAGGACTCCATCAGATACTATTCTCTTGAAATAGTATCTGATGGAGTCCGGGGA-3’) between MluI and ClaI restriction sites in the pLVTHM plasmid. Transfection of HEK-293 T cells (kindly provided by Genethon, Evry, France) with pLV-tTR-KRAB-Red or pLVTHM, and preparation of high titer lentiviruses pseudotyped with VSV-G protein have been performed as previously described (82). pLV-tTRKRAB-red and pLVTHM were a gift from Didier Trono (Addgene plasmid # 12250; http://n2t.net/addgene:12250; RRID:Addgene_12250 and Addgene plasmid # 12247; http://n2t.net/addgene:12247; RRID:Addgene_12247) (83).

### RNA interferences

HeLa cells were seeded at 250.000 cells per well in a 6-wells plate. siRNAs oligonucleotides (Table) were transfected twice (24 and 48 hours after seeding) at 50nM final concentration per well with Lipofectamine RNAiMax Reagent (Invitrogen) according to manufacturer’s recommendations. For TOP2A and TOP2B co-depletion, each siRNA was used at the final concentration of 25 nM. Cells were split 24h after the second-round transfection for immunodetection, immunoblotting and viability assays, and were treated 24h after being seeded.

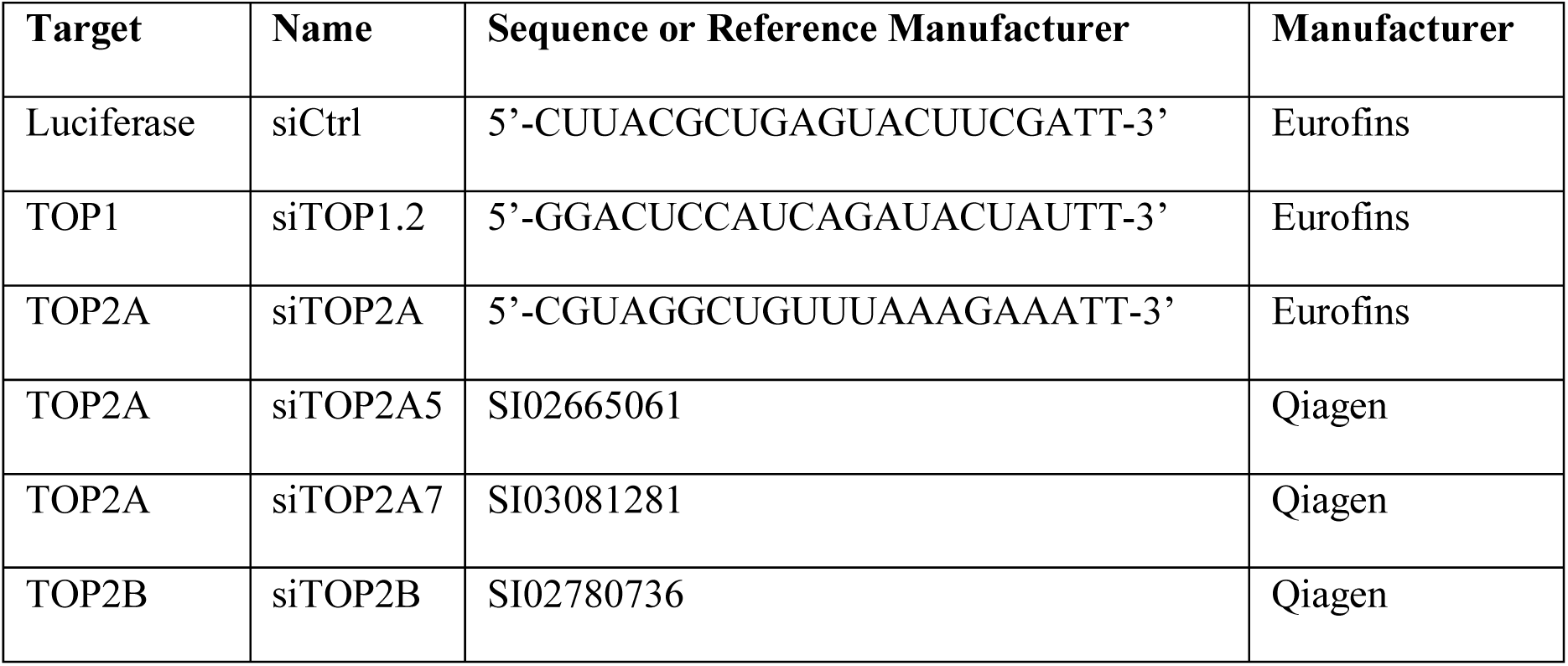

### RNA-seq

RNAseq was performed at the GeT-PlaGe core facility, INRA Toulouse from total RNA prepared with the RNeasy Plus Mini Kit (Qiagen) according to the manufacturer’s instructions. RNA-seq libraries were prepared according to Illumina’s protocols using the Illumina TruSeq Stranded mRNA sample prep kit. Briefly, mRNAs were selected using poly-dT beads. Then, RNAs were fragmented and adaptators ligated. Eleven cycles of PCR were applied for libraries amplification. Library quality was assessed using a Fragment Analyser System (Agilent) and libraries were quantified by Q-PCR using the Kapa Library Quantification Kit (Roche). RNA-seq experiments were performed on an Illumina HiSeq3000 using a paired-end read length of 2×150 pb.

### RNA-Seq alignment and SNP prediction and filtering

Read quality was checked within the ng6 environment (84) using fastQC (http://www.bioinformatics.babraham.ac.uk/projects/fastqc/) and Burrows-Wheeler Aligner BWA (85) to search for contamination. The reads were cleaned with cutadapt v 1.8.3 and aligned against hg38 reference human genome with STAR v2.5.2b (86). Expression levels were computed with featureCount (87) using Ensembl annotation. Alignements were deduplicated with samtools rmdup and reads not uniquely mapped removed. Then GATK v3.5 base quality score recalibration was applied (88). Indel realignment, SNP and INDEL discovery were performed with HaplotypeCaller using standard hard filtering parameters according to GATK Best Practices recommendations for RNAseq. Finally variants were annotated using snpEff v4.3T (89). A python script was used to select protein coding variants specific to CXR clones as compared to wild-type HAP1, with a minimal allele frequency of 0.9 and a depth greater than 10 reads. Among these variants, we selected variants resulting in frameshifts, mis- and non-sense mutations as compared to the reference human genome hg38. Cytoscape v3.2.0 (90) was used to identify genes found mutated in several CXR clones. Upon TOP2A identification as a common gene mutated in 5 CXR clones, IGV v2.4.15 was used to scrutinize alignment data and revealed two TOP2A mutations missed by the analysis: for CXR#A2, the point mutation S654I and, for CXR#A6, a mutation of the first nucleotide of the last intron leading to intron retention. Clone clustering under Cytoscape based on shared mutated genes suggested a common origin for clones CXR#A1, #A3, #A5 and #B4 (multiple common mutations).

### Targeted sequencing of TOP2A cDNA from HAP1 clones

Total RNAs were extracted from wild-type or F14R HAP1 with the RNeasy Plus Mini Kit (Qiagen) according to the manufacturer’s instructions. TOP2A cDNA was produced from these RNAs with the Superscript III First-Strand kit (Thermo Fisher Scientific) according to the manufacturer’s instructions and using the TOP2A-Rv primer. The resulting TOP2A cDNAs was amplified in 4 overlapping fragments using the primer pairs [TOP2A-F1, TOP2A-R1], [TOP2A-F2, TOP2A-R2], [TOP2A-F3, TOP2A-R3] and [TOP2A-F4, TOP2A-Rv] and sequenced using the same primers except for the last fragment for which the TOP2A-R4 sequencing primer was also used.

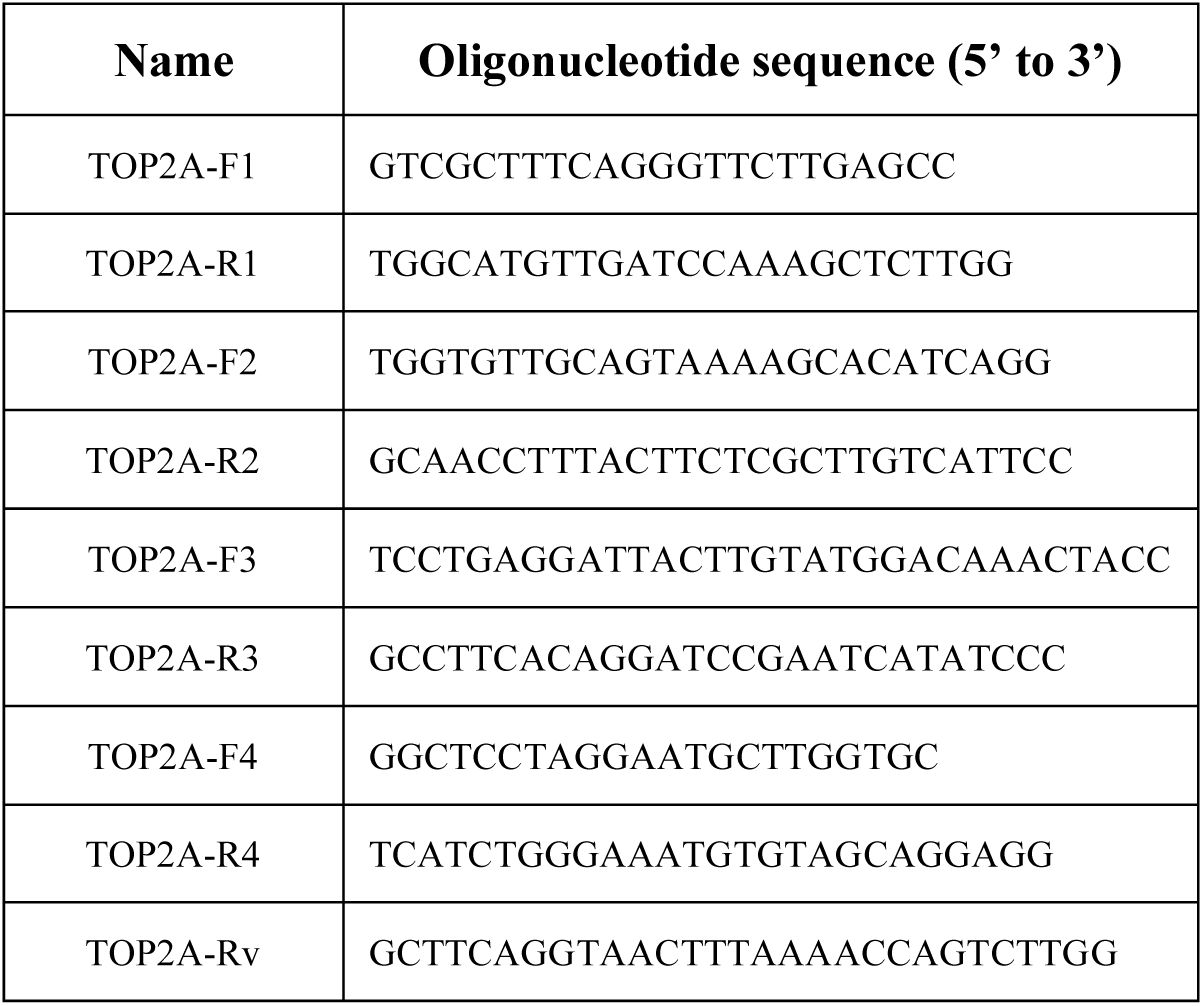

### Heparin-based extraction for TOP2cc immunodetection

This method was adapted from (48). Briefly, HAP1 cells were seeded at 1.5 10^6^ cells in 6 cm-dishes 24 hours prior etoposide treatment (200 µM, 1h). After treatment, cells were harvested with trypsin and washed with cold PBS. After gentle centrifugation, cells were resuspended in lysis buffer (150mM NaCl, 1mM EDTA, 0.5X NP40, 2X HALT Protease and Phosphatase Inhibitor Cocktail (ThermoScientific) 20mM Tris-HCl, pH8) complemented with 100U/mL Heparin (Sigma) and incubated on ice for 15 min. Then lysates were centrifugated at 15000 rpm at 4°C for 5min and pellets were resuspended with lysis buffer. In order to facilitate migration on polyacrylamide gel, a sonication was performed to degrade DNA present within the extracts. Protein concentrations were determined by measuring absorbance at 280nm (Nanodrop) and heparin-based extracts were diluted with denaturing lysis buffer (4% SDS, 20% glycerol and 120 mM Tris-HCl, pH 6.8). Western blot was performed as described below.

### Cell lysis and western blotting

Whole-cell extracts were prepared from PBS-washed pellet lysed with denaturing lysis buffer (4% SDS, 20% glycerol and 120 mM Tris-HCl, pH 6.8) and 10 strokes through a 24G needle. Protein concentration were determined by measuring absorbance at 280nm (Nanodrop). For loading, an equal volume of a solution of 0.01% bromophenol blue and 200 mM dithiothreitol was added to the extracts then boiled at 95°C for 5 min. About 80 µg of denatured proteins were loaded for each condition and separated on gradient 4–12% polyacrylamide classic or TGX Stain-Free pre-cast gels (Biorad) and transferred onto nitrocellulose membrane (Biorad). Before blocking (0.1% tween20, non-fat dry milk 5% and PBS), ponceau S staining or UV exposition of membrane (for Stain Free gels) was used to confirm homogeneous loading. The membrane was successively probed with primary antibody and appropriate goat secondary antibodies coupled to horseradish peroxidase (described in table below). Chemidoc imager (Biorad) was used to perform UV and Clarity ECL (Biorad) detection. Digital data were processed and quantified using ImageJ software.

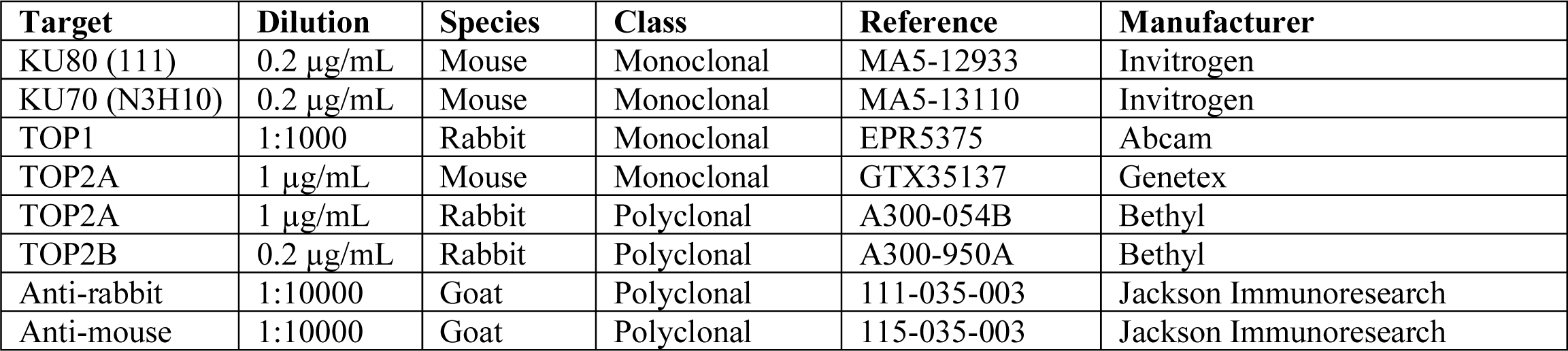

| Target | Dilution | Species | Class | Reference | Manufacturer |
| --- | --- | --- | --- | --- | --- |
| KU80 (111) | 0.2 µg/mL | Mouse | Monoclonal | MA5-12933 | Invitrogen |
| KU70 (N3H10) | 0.2 µg/mL | Mouse | Monoclonal | MA5-13110 | Invitrogen |
| TOP1 | 1:1000 | Rabbit | Monoclonal | EPR5375 | Abcam |
| TOP2A | 1 µg/mL | Mouse | Monoclonal | GTX35137 | Genetex |
| TOP2A | 1 µg/mL | Rabbit | Polyclonal | A300-054B | Bethyl |
| TOP2B | 0.2 µg/mL | Rabbit | Polyclonal | A300-950A | Bethyl |
| Anti-rabbit | 1:10000 | Goat | Polyclonal | 111-035-003 | Jackson ImmunoResearch |
| Anti-mouse | 1:10000 | Goat | Polyclonal | 115-035-003 | Jackson ImmunoResearch |

### Immunofluorescence

RPE1-hTERT cells were seeded, treated and stained in the same conditions than HeLa cells. HeLa cells and HAP1 cells were seeded in 24-wells plate at respectively 100.000 cells/well and 25.000 cells/well on glass coverslips (VWR, #631-0150). HeLa cells and HAP1 cells were respectively treated 24 hours and 48 hours later, and then fixed with paraformaldehyde 2% in PBS at room temperature (10 min for HeLa cells, 15 min for HAP1 cells), washed with PBS and permeabilized for 15 min at room temperature with 10 mM Tris–HCl pH 7.5, 120 mM KCl, 20 mM NaCl, 0.1% Triton-X 100. In EdU treated cells, cells were washed with PBS and EdU detection reaction was performed with Click-iT RNA Alexa Fluor Imaging Kit according to manufacturer’s recommendation, with 2uM Alexa Fluor 594 or 648 azide for 30 min at room temperature. Then, cells were washed with PBS and incubated for about 1 h at 37°C in blocking buffer (20 mM Tris–HCl pH 7.5, 150 mM NaCl, 2% BSA, 0.2% fish gelatin, 0.1% Triton-X 100) prior to incubation overnight at 4°C with primary antibody diluted in blocking buffer (dilutions shown in table below). For BG4 immunodetection, blocking buffer were complemented 0.3 µg/µl of RNAse A (91). Cells were then washed with PBS-Tween20 0.1% and incubated with appropriate secondary goat antibody coupled to AlexaFluor 488 or 594 diluted in blocking buffer (dilutions shown in table below) for 1 h at room temperature. At last, cells were washed with PBS-Tween20 0.1% and stained with 0.1 μg/mL DAPI for 20 min at room temperature, and coverslips were mounted with Vectashield mounting medium (Vector Laboratories). Nuclear γH2AX foci, 53BP1 foci, BG4 foci and EdU integrated density staining overlapping with DAPI staining were quantified with ImageJ software. Nuclear DAPI integrated density staining was quantified with ImageJ software and correlated to nuclear EdU integrated density to determine cell cycle phase for each cells as described in (92). Quantifications of nuclear γH2AX and 53BP1 foci induced by G4 ligands are represented normalized to non-treated (NT) conditions.

| Target | Dilution | Species | Class | Reference | Manufacturer |
| --- | --- | --- | --- | --- | --- |
| 53BP1 | 1.3 μg/mL | Mouse | Monoclonal | MAB-3803 | Millipore |
| γH2AX (Phospho S139) | 0.7 μg/mL | Rabbit | Monoclonal | 81299 | Abcam |
| BG4 | 0.25 μg/mL | Mouse | Monoclonal | Ab00174-1.1 | Absolute antibody |
| TOP2A | 0.2 μg/mL | Mouse | Monoclonal | GTX35137 | Genetex |
| Anti-rabbit 488 | 2 μg/mL | Goat | Polyclonal | A11008 | Invitrogen |
| Anti-mouse 594 | 2 μg/mL | Goat | Polyclonal | A11005 | Invitrogen |
| Anti-mouse 488 | 2 μg/mL | Goat | Polyclonal | A11001 | Invitrogen |
| Anti-rabbit 594 | 2 μg/mL | Goat | Polyclonal | A11012 | Invitrogen |

### Viability assay (SRB)

HAP1 cells were seeded in 96-flat-wells plate at 3500 cells per well. Serial dilutions of various compounds were realized allowing same solvent concentration for each condition, and cells were treated 24 hours after seeding. After 3 days, HAP1 cells were fixed with 10% trichloroacetic acid for 1h at 4°C, washed and dried overnight. Protein content of cells were stained by 0.057% sulforhodamin B in 1% acetic acid for 30 min at room temperature, then cells were washed with 1% acetic acid and dried overnight. Finally, 200 μl of a 10 mM Tris-base solution was added, plates were agitated for 1h at room temperature and SRB levels were measured by absorbance at 490 nm using μQuant microplate spectrophotometer (Bio-Tek Instruments). Percentages of cell viability are expressed after normalization relative to non-treated controls. For characterization of drug-resistant HAP1, the curve fitting analysis (nonlinear regression curve fit) was performed with the algorithm provided by the GraphPad Software (version 8) and allowed to calculate the mean IC_50_ value (50% inhibitory concentration) under each condition.

### Clonogenic assay

Clonogenic assay was performed as described by (49). Briefly, after transfection with siRNA, HeLa cells were seeded at low density (250 cells/well) the day before treatment, pre-incubated with 5,6-dichloro-1-beta-D-ribofuranosylbenzimidazole (DRB) or dimethylsulfoxide (DMSO) for 1 h and treated for 4 hours with pyridostatin in presence of transcription inhibitor or DMSO before being removed and fresh medium was added. After 10–15 days, cells were stained with crystal violet and the colonies were counted (at least, 50 colonies were counted for each condition per experiment). Data were normalized to the non-treated conditions. The curve fitting analysis (nonlinear regression curve fit) was performed with the algorithm provided by the GraphPad Software (version 8) and allowed to calculate the mean IC50 value (50% inhibitory concentration) under each condition.

### Statistical analyses

All results provide from at least three independent experiments. Statistical analyses were performed with GraphPad Prism Software (version 8). For γH2AX and 53BP1 quantifications analyses, multiple unpaired t-tests (without corrections for multiple comparisons) were performed between pairs of conditions. For BG4 quantifications analyses, results of at least three independent experiments were pulled together and unpaired Welch’s t-tests were performed between pairs of conditions. On all figures, significant differences between specified pairs of conditions are shown by asterisks (*: p-value<0.05; **: p-value<0.01; ***: p-value<0.0005; ****: p-value<0.0001). NS is for non-significant difference.

## References

1. A. Aguilera, The connection between transcription and genomic instability. EMBO J 21, 195–201 (2002).

2. H. Gaillard, E. Herrera-Moyano, A. Aguilera, Transcription-associated genome instability. Chem Rev 113, 8638–8661 (2013).

3. B. Gomez-Gonzalez, A. Aguilera, Transcription-mediated replication hindrance: a major driver of genome instability. Genes Dev 33, 1008–1026 (2019).

4. A. Aguilera, T. Garcia-Muse, R loops: from transcription byproducts to threats to genome stability. Mol Cell 46, 115–124 (2012).

5. A. De Magis et al., DNA damage and genome instability by G-quadruplex ligands are mediated by R loops in human cancer cells. Proc Natl Acad Sci U S A 116, 816–825 (2019).

6. S. Burge, G. N. Parkinson, P. Hazel, A. K. Todd, S. Neidle, Quadruplex DNA: sequence, topology and structure. Nucleic Acids Res 34, 5402–5415 (2006).

7. V. S. Chambers et al., High-throughput sequencing of DNA G-quadruplex structures in the human genome. Nat Biotechnol 33, 877–881 (2015).

8. R. Hansel-Hertsch et al., G-quadruplex structures mark human regulatory chromatin. Nat Genet 48, 1267–1272 (2016).

9. R. Hansel-Hertsch, J. Spiegel, G. Marsico, D. Tannahill, S. Balasubramanian, Genome-wide mapping of endogenous G-quadruplex DNA structures by chromatin immunoprecipitation and high-throughput sequencing. Nat Protoc 13, 551–564 (2018).

10. R. Hansel-Hertsch, M. Di Antonio, S. Balasubramanian, DNA G-quadruplexes in the human genome: detection, functions and therapeutic potential. Nat Rev Mol Cell Biol 18, 279–284 (2017).

11. P. Murat, S. Balasubramanian, Existence and consequences of G-quadruplex structures in DNA. Curr Opin Genet Dev 25, 22–29 (2014).

12. N. Maizels, G4-associated human diseases. EMBO Rep 16, 910–922 (2015).

13. L. K. Lerner, J. E. Sale, Replication of G Quadruplex DNA. Genes (Basel) 10 (2019).

14. R. Rodriguez et al., Small-molecule-induced DNA damage identifies alternative DNA structures in human genes. Nat Chem Biol 8, 301–310 (2012).

15. N. van Wietmarschen et al., BLM helicase suppresses recombination at G-quadruplex motifs in transcribed genes. Nat Commun 9, 271 (2018).

16. N. Kim, The Interplay between G-quadruplex and Transcription. Curr Med Chem 26, 2898–2917 (2019).

17. N. Puget, K. M. Miller, G. Legube, Non-canonical DNA/RNA structures during Transcription-Coupled Double-Strand Break Repair: Roadblocks or Bona fide repair intermediates? DNA Repair (Amst) 81, 102661 (2019).

18. L. Chen et al., R-ChIP Using Inactive RNase H Reveals Dynamic Coupling of R-loops with Transcriptional Pausing at Gene Promoters. Mol Cell 68, 745–757 e745 (2017).

19. J. Eddy et al., G4 motifs correlate with promoter-proximal transcriptional pausing in human genes. Nucleic Acids Res 39, 4975–4983 (2011).

20. Y. Pommier, Y. Sun, S. N. Huang, J. L. Nitiss, Roles of eukaryotic topoisomerases in transcription, replication and genomic stability. Nat Rev Mol Cell Biol 17, 703–721 (2016).

21. P. Yadav, N. Owiti, N. Kim, The role of topoisomerase I in suppressing genome instability associated with a highly transcribed guanine-rich sequence is not restricted to preventing RNA:DNA hybrid accumulation. Nucleic Acids Res 44, 718–729 (2016).

22. M. Drolet et al., The problem of hypernegative supercoiling and R-loop formation in transcription. Front Biosci 8, d210–221 (2003).

23. A. El Hage, S. L. French, A. L. Beyer, D. Tollervey, Loss of Topoisomerase I leads to R-loop-mediated transcriptional blocks during ribosomal RNA synthesis. Genes Dev 24, 1546–1558 (2010).

24. S. G. Manzo et al., DNA Topoisomerase I differentially modulates R-loops across the human genome. Genome Biol 19, 100 (2018).

25. P. Yadav et al., Topoisomerase I plays a critical role in suppressing genome instability at a highly transcribed G-quadruplex-forming sequence. PLoS Genet 10, e1004839 (2014).

26. J. L. Nitiss, DNA topoisomerase II and its growing repertoire of biological functions. Nat Rev Cancer 9, 327–337 (2009).

27. R. Madabhushi, The Roles of DNA Topoisomerase IIbeta in Transcription. Int J Mol Sci 19 (2018).

28. J. E. Deweese, N. Osheroff, The DNA cleavage reaction of topoisomerase II: wolf in sheep’s clothing. Nucleic Acids Res 37, 738–748 (2009).

29. J. L. Nitiss, Targeting DNA topoisomerase II in cancer chemotherapy. Nat Rev Cancer 9, 338–350 (2009).

30. Y. Pommier, E. Leo, H. Zhang, C. Marchand, DNA topoisomerases and their poisoning by anticancer and antibacterial drugs. Chem Biol 17, 421–433 (2010).

31. J. R. Fan et al., Cellular processing pathways contribute to the activation of etoposide-induced DNA damage responses. DNA Repair (Amst) 7, 452–463 (2008).

32. Y. Mao, S. D. Desai, C. Y. Ting, J. Hwang, L. F. Liu, 26 S proteasome-mediated degradation of topoisomerase II cleavable complexes. J Biol Chem 276, 40652–40658 (2001).

33. A. Zhang et al., A protease pathway for the repair of topoisomerase II-DNA covalent complexes. J Biol Chem 281, 35997–36003 (2006).

34. T. Aparicio, R. Baer, M. Gottesman, J. Gautier, MRN, CtIP, and BRCA1 mediate repair of topoisomerase II-DNA adducts. J Cell Biol 212, 399–408 (2016).

35. K. Nakamura et al., Collaborative action of Brca1 and CtIP in elimination of covalent modifications from double-strand breaks to facilitate subsequent break repair. PLoS Genet 6, e1000828 (2010).

36. F. Gomez-Herreros et al., TDP2-dependent non-homologous end-joining protects against topoisomerase II-induced DNA breaks and genome instability in cells and in vivo. PLoS Genet 9, e1003226 (2013).

37. M. E. Ashour, R. Atteya, S. F. El-Khamisy, Topoisomerase-mediated chromosomal break repair: an emerging player in many games. Nat Rev Cancer 15, 137–151 (2015).

38. A. Khot et al., First-in-Human RNA Polymerase I Transcription Inhibitor CX-5461 in Patients with Advanced Hematologic Cancers: Results of a Phase I Dose-Escalation Study. Cancer Discov 9, 1036–1049 (2019).

39. M. Haddach, et al., Discovery of CX-5461, the First Direct and Selective Inhibitor of RNA Polymerase I, for Cancer Therapeutics. ACS Med Chem Lett 3, 602–606 (2012).

40. S. S. Negi, P. Brown, Transient rRNA synthesis inhibition with CX-5461 is sufficient to elicit growth arrest and cell death in acute lymphoblastic leukemia cells. Oncotarget 6, 34846–34858 (2015).

41. H. Xu, et al., CX-5461 is a DNA G-quadruplex stabilizer with selective lethality in BRCA1/2 deficient tumours. Nat Commun 8, 14432 (2017).

42. J. V. Forment et al., Genome-wide genetic screening with chemically mutagenized haploid embryonic stem cells. Nat Chem Biol 13, 12–14 (2017).

43. C. Kasap, O. Elemento, T. M. Kapoor, DrugTargetSeqR: a genomics- and CRISPR-Cas9-based method to analyze drug targets. Nat Chem Biol 10, 626–628 (2014).

44. S. A. Wacker, B. R. Houghtaling, O. Elemento, T. M. Kapoor, Using transcriptome sequencing to identify mechanisms of drug action and resistance. Nat Chem Biol 8, 235–237 (2012).

45. N. Akimitsu et al., Enforced cytokinesis without complete nuclear division in embryonic cells depleting the activity of DNA topoisomerase IIalpha. Genes Cells 8, 393–402 (2003).

46. A. J. Carpenter, A. C. Porter, Construction, characterization, and complementation of a conditional-lethal DNA topoisomerase IIalpha mutant human cell line. Mol Biol Cell 15, 5700–5711 (2004).

47. J. V. Walker, J. L. Nitiss, DNA topoisomerase II as a target for cancer chemotherapy. Cancer Invest 20, 570–589 (2002).

48. M. de Campos-Nebel, M. Palmitelli, M. Gonzalez-Cid, A flow cytometry-based method for a high-throughput analysis of drug-stabilized topoisomerase II cleavage complexes in human cells. Cytometry A 89, 852–860 (2016).

49. O. Bombarde et al., The DNA-Binding Polyamine Moiety in the Vectorized DNA Topoisomerase II Inhibitor F14512 Alters Reparability of the Consequent Enzyme-Linked DNA Double-Strand Breaks. Mol Cancer Ther 16, 2166–2177 (2017).

50. R. Rodriguez et al., A novel small molecule that alters shelterin integrity and triggers a DNA-damage response at telomeres. J Am Chem Soc 130, 15758–15759 (2008).

51. N. R. Pannunzio, G. Watanabe, M. R. Lieber, Nonhomologous DNA end-joining for repair of DNA double-strand breaks. J Biol Chem 293, 10512–10523 (2018).

52. M. Kawatani et al., Identification of a small-molecule inhibitor of DNA topoisomerase II by proteomic profiling. Chem Biol 18, 743–751 (2011).

53. K. Yankulov, K. Yamashita, R. Roy, J. M. Egly, D. L. Bentley, The transcriptional elongation inhibitor 5,6-dichloro-1-beta-D-ribofuranosylbenzimidazole inhibits transcription factor IIH-associated protein kinase. J Biol Chem 270, 23922–23925 (1995).

54. S. Britton et al., DNA damage triggers SAF-A and RNA biogenesis factors exclusion from chromatin coupled to R-loops removal. Nucleic Acids Res 42, 9047–9062 (2014).

55. J. J. Champoux, DNA topoisomerases: structure, function, and mechanism. Annu Rev Biochem 70, 369–413 (2001).

56. A. C. Cheung, P. Cramer, A movie of RNA polymerase II transcription. Cell 149, 1431–1437 (2012).

57. K. W. Zheng et al., Superhelicity Constrains a Localized and R-Loop-Dependent Formation of G-Quadruplexes at the Upstream Region of Transcription. ACS Chem Biol 12, 2609–2618 (2017).

58. M. R. Gartenberg, J. C. Wang, Positive supercoiling of DNA greatly diminishes mRNA synthesis in yeast. Proc Natl Acad Sci U S A 89, 11461–11465 (1992).

59. R. S. Joshi, B. Pina, J. Roca, Positional dependence of transcriptional inhibition by DNA torsional stress in yeast chromosomes. EMBO J 29, 740–748 (2010).

60. L. Baranello et al., RNA Polymerase II Regulates Topoisomerase 1 Activity to Favor Efficient Transcription. Cell 165, 357–371 (2016).

61. H. Bunch et al., Transcriptional elongation requires DNA break-induced signalling. Nat Commun 6, 10191 (2015).

62. G. I. Dellino et al., Release of paused RNA polymerase II at specific loci favors DNA double-strand-break formation and promotes cancer translocations. Nat Genet 51, 1011–1023 (2019).

63. D. A. Scicchitano, I. Mellon, Transcription and DNA damage: a link to a kink. Environ Health Perspect 105 Suppl 1, 145–153 (1997).

64. S. Singh et al., Pausing sites of RNA polymerase II on actively transcribed genes are enriched with DNA double-stranded breaks. J Biol Chem 10.1074/jbc.RA119.011665 (2020).

65. T. R. Blower et al., A complex suite of loci and elements in eukaryotic type II topoisomerases determine selective sensitivity to distinct poisoning agents. Nucleic Acids Res 47, 8163–8179 (2019).

66. J. P. McPherson, G. J. Goldenberg, Induction of apoptosis by deregulated expression of DNA topoisomerase IIalpha. Cancer Res 58, 4519–4524 (1998).

67. S. Ray et al., Topoisomerase IIalpha promotes activation of RNA polymerase I transcription by facilitating pre-initiation complex formation. Nat Commun 4, 1598 (2013).

68. K. I. McLuckie et al., G-quadruplex DNA as a molecular target for induced synthetic lethality in cancer cells. J Am Chem Soc 135, 9640–9643 (2013).

69. E. Salvati et al., Lead Discovery of Dual G-Quadruplex Stabilizers and Poly(ADP-ribose) Polymerases (PARPs) Inhibitors: A New Avenue in Anticancer Treatment. J Med Chem 60, 3626–3635 (2017).

70. C. A. Austin et al., TOP2B: The First Thirty Years. Int J Mol Sci 19 (2018).

71. N. Mondal, J. D. Parvin, DNA topoisomerase IIalpha is required for RNA polymerase II transcription on chromatin templates. Nature 413, 435–438 (2001).

72. M. Tammaro, P. Barr, B. Ricci, H. Yan, Replication-dependent and transcription-dependent mechanisms of DNA double-strand break induction by the topoisomerase 2-targeting drug etoposide. PLoS One 8, e79202 (2013).

73. N. Kim, S. Jinks-Robertson, The Top1 paradox: Friend and foe of the eukaryotic genome. DNA Repair (Amst) 56, 33–41 (2017).

74. R. N. Ganapathi, M. K. Ganapathi, Mechanisms regulating resistance to inhibitors of topoisomerase II. Front Pharmacol 4, 89 (2013).

75. M. Y. Kim, W. Duan, M. Gleason-Guzman, L. H. Hurley, Design, synthesis, and biological evaluation of a series of fluoroquinoanthroxazines with contrasting dual mechanisms of action against topoisomerase II and G-quadruplexes. J Med Chem 46, 571–583 (2003).

76. G. Zoidis et al., Indenocinnoline derivatives as G-quadruplex binders, topoisomerase IIalpha inhibitors and antiproliferative agents. Bioorg Med Chem 25, 2625–2634 (2017).

77. S. Raje, R. Barthwal, Molecular recognition of 3+1 hybrid human telomeric G-quadruplex DNA d-[AGGG(TTAGGG)3] by anticancer drugs epirubicin and adriamycin leads to thermal stabilization. Int J Biol Macromol 139, 1272–1287 (2019).

78. S. Raje, K. Pandav, R. Barthwal, Dual mode of binding of anti cancer drug epirubicin to G-quadruplex [d-(TTAGGGT)]4 containing human telomeric DNA sequence induces thermal stabilization. Bioorg Med Chem 27, 115131 (2019).

79. S. Raje, K. Pandav, R. Barthwal, Binding of anticancer drug adriamycin to parallel G-quadruplex DNA [d-(TTAGGGT)]4 comprising human telomeric DNA leads to thermal stabilization: A multiple spectroscopy study. J Mol Recognit 33, e2815 (2020).

80. S. Tripathi, R. Barthwal, NMR based structure reveals groove binding of mitoxantrone to two sites of [d-(TTAGGGT)]4 having human telomeric DNA sequence leading to thermal stabilization of G-quadruplex. Int J Biol Macromol 111, 326–341 (2018).

81. P. M. Bruno et al., The primary mechanism of cytotoxicity of the chemotherapeutic agent CX-5461 is topoisomerase II poisoning. Proc Natl Acad Sci U S A 10.1073/pnas.1921649117 (2020).

82. C. Delenda, Lentiviral vectors: optimization of packaging, transduction and gene expression. J Gene Med 6 Suppl 1, S125–138 (2004).

83. M. Wiznerowicz, D. Trono, Conditional suppression of cellular genes: lentivirus vector-mediated drug-inducible RNA interference. J Virol 77, 8957–8961 (2003).

84. J. Mariette et al., NG6: Integrated next generation sequencing storage and processing environment. BMC Genomics 13, 462 (2012).

85. H. Li, R. Durbin, Fast and accurate short read alignment with Burrows-Wheeler transform. Bioinformatics 25, 1754–1760 (2009).

86. A. Dobin et al., STAR: ultrafast universal RNA-seq aligner. Bioinformatics 29, 15–21 (2013).

87. Y. Liao, G. K. Smyth, W. Shi, featureCounts: an efficient general purpose program for assigning sequence reads to genomic features. Bioinformatics 30, 923–930 (2014).

88. A. McKenna et al., The Genome Analysis Toolkit: a MapReduce framework for analyzing next-generation DNA sequencing data. Genome Res 20, 1297–1303 (2010).

89. P. Cingolani et al., A program for annotating and predicting the effects of single nucleotide polymorphisms, SnpEff: SNPs in the genome of Drosophila melanogaster strain w1118; iso-2; iso-3. Fly (Austin) 6, 80–92 (2012).

90. P. Shannon et al., Cytoscape: a software environment for integrated models of biomolecular interaction networks. Genome Res 13, 2498–2504 (2003).

91. A. P. David et al., CNBP controls transcription by unfolding DNA G-quadruplex structures. Nucleic Acids Res 47, 7901–7913 (2019).

92. V. Roukos, G. Pegoraro, T. C. Voss, T. Misteli, Cell cycle staging of individual cells by fluorescence microscopy. Nat Protoc 10, 334–348 (2015).

