## Supplementary Figures and Legends for "Transcription-associated topoisomerase activities control DNA-breaks production by G-quadruplex ligands"

| | Nocodazole | | Etoposide | | F14512 | | CX5461 | | PDS | |
|---|---|---|---|---|---|---|---|---|---|---|
| | $IC_{50}$ value (µM) | RI* | $IC_{50}$ value (µM) | RI* | $IC_{50}$ value (nM) | RI* | $IC_{50}$ value (µM) | RI* | $IC_{50}$ value (µM) | RI* |
| HAP1 WT | 0.02 | | 0.05 | | nd | | 0.03 | | 4.20 | |
| CXR #A1 | 0.02 | 1.10 | 0.39 | 8.15 | nd | nd | 0.29 | 8.91 | 17.09 | 4.07 |
| CXR #A2 | 0.02 | 0.90 | 0.72 | 14.92 | nd | nd | 0.27 | 8.30 | 11.50 | 2.74 |
| CXR #A3 | 0.02 | 0.90 | 0.45 | 9.35 | nd | nd | 0.22 | 6.76 | 14.94 | 3.56 |
| CXR #A5 | 0.02 | 0.90 | 0.52 | 10.79 | nd | nd | 0.26 | 7.94 | 18.27 | 4.35 |
| CXR #A6 | 0.02 | 0.90 | 0.64 | 13.42 | nd | nd | 0.26 | 8.00 | 15.63 | 3.72 |
| CXR #B3 | 0.02 | 1.00 | 0.91 | 19.02 | nd | nd | 0.34 | 10.15 | 17.57 | 4.19 |
| CXR #B4 | 0.02 | 0.90 | 0.51 | 10.67 | nd | nd | 0.26 | 7.82 | 18.41 | 4.39 |
| HAP1 WT | 0.02 | | 0.05 | | 2.73 | | 0.04 | | 4.16 | |
| F14R #A1 | 0.02 | 0.90 | 0.15 | 2.68 | 23.92 | 8.75 | 0.17 | 4.71 | 15.23 | 3.67 |
| F14R #A2 | 0.02 | 1.23 | 0.27 | 5.01 | 40.83 | 14.93 | 0.33 | 9.15 | 12.94 | 3.11 |
| F14R #A3 | 0.02 | 1.20 | 0.22 | 4.11 | 28.82 | 10.54 | 0.20 | 5.64 | 10.95 | 2.64 |
| F14R #A5 | 0.02 | 1.20 | 0.29 | 5.36 | 42.07 | 15.39 | 0.35 | 9.71 | 14.74 | 3.55 |
| F14R #A6 | 0.02 | 1.32 | 0.42 | 7.69 | 29.95 | 10.95 | 0.13 | 3.58 | 7.76 | 1.87 |
| F14R #C1 | 0.02 | 1.13 | 0.35 | 6.38 | 42.50 | 15.54 | 0.60 | 16.82 | 17.37 | 4.18 |
| F14R #C5 | 0.01 | 0.65 | 0.15 | 2.76 | 28.14 | 10.29 | 0.27 | 7.57 | 16.24 | 3.91 |
| F14R #C6 | 0.02 | 0.99 | 0.18 | 3.23 | 33.20 | 12.14 | 0.34 | 9.55 | 8.41 | 2.02 |

*RI: Resistance Index; ratio $\frac{IC_{50}\,(CXR)\ or\ (F14R)}{IC_{50}(HAP1\,WT)}$

**Supplementary Table 1:** $IC_{50}$ values as assessed by cell survival assays of nocodazole, etoposide and G4 ligands (CX5461 and PDS) on WT CX5461 (CXR) and F14512 resistant (F14R) HAP1 cells.

| CXR#A1 | Mutation (Nt) | Mutation (aa) | CXR#A5 | | |
|---|---|---|---|---|---|
| BPTF | c.1775A>G | p.Lys592Arg | BPTF | c.1775A>G | p.Lys592Arg |
| DOK4 | c.514C>T | p.Pro172Ser | DOK4 | c.514C>T | p.Pro172Ser |
| EIF2B1 | c.778G>A | p.Ala260Thr | EIF2B1 | c.778G>A | p.Ala260Thr |
| KCTD5 | c.586G>T | p.Glu196* | KCTD5 | c.586G>T | p.Glu196* |
| KIF13A | c.3982G>T | p.Ala1328Ser | LATS2 | c.2732C>T | p.Pro911Leu |
| LATS2 | c.2732C>T | p.Pro911Leu | RBM26 | c.749C>T | p.Thr250Ile |
| RBM26 | c.749C>T | p.Thr250Ile | SPRED2 | c.1232G>T | p.Gly411Val |
| SPRED2 | c.1232G>T | p.Gly411Val | SPRED2 | c.1231G>T | p.Gly411Cys |
| SPRED2 | c.1231G>T | p.Gly411Cys | TOP2A | c.253T>A | p.Phe85Ile |
| TOP2A | c.253T>A | p.Phe85Ile | | | |
| | | | **CXR#A6** | | |
| **CXR#A2** | | | G3BP1 | c.524A>G | p.Asp175Gly |
| ADCK2 | c.1357G>T | p.Val453Phe | SULF2 | c.1946G>A | p.Arg649Gln |
| ARHGEF12 | c.4335C>G | p.Ile1445Met | | | |
| B4GALNT4 | c.2613G>T | p.Glu871Asp | **CXR#B3** | | |
| CALR | c.301C>T | p.Gln101* | ETNK1 | c.229G>T | p.Ala77Ser |
| JDP2 | c.515A>T | p.Glu172Val | FAM219A | c.211G>A | p.Gly71Ser |
| KIAA1211 | c.3314C>A | p.Ala1105Glu | GLIPR2 | c.122T>A | p.Val41Asp |
| TOX2 | c.498C>A | p.Ser166Arg | KDM5B | c.1481G>T | p.Gly494Val |
| | | | KDM5B | c.1480G>T | p.Gly494Cys |
| **CXR#A3** | | | RAF1 | c.976G>T | p.Glu326* |
| BPTF | c.1775A>G | p.Lys592Arg | TBC1D24 | c.637C>A | p.Gln213Lys |
| DOK4 | c.514C>T | p.Pro172Ser | TESMIN | c.596C>A | p.Ser199Tyr |
| EIF2B1 | c.778G>A | p.Ala260Thr | TOP2A | c.2107C>A | p.Leu703Ile |
| KCTD5 | c.586G>T | p.Glu196* | | | |
| KIF13A | c.3982G>T | p.Ala1328Ser | **CXR#B4** | | |
| LATS2 | c.2732C>T | p.Pro911Leu | BPTF | c.1775A>G | p.Lys592Arg |
| RBM26 | c.749C>T | p.Thr250Ile | DOK4 | c.514C>T | p.Pro172Ser |
| SPRED2 | c.1232G>T | p.Gly411Val | EIF2B1 | c.778G>A | p.Ala260Thr |
| SPRED2 | c.1231G>T | p.Gly411Cys | KCTD5 | c.586G>T | p.Glu196* |
| TOP2A | c.253T>A | p.Phe85Ile | LATS2 | c.2732C>T | p.Pro911Leu |
| | | | RBM26 | c.749C>T | p.Thr250Ile |
| | | | RREB1 | c.2567G>T | p.Cys856Phe |
| | | | SPRED2 | c.1232G>T | p.Gly411Val |
| | | | SPRED2 | c.1231G>T | p.Gly411Cys |
| | | | TOP2A | c.253T>A | p.Phe85Ile |

**Supplementary table 2:** non- or false-sense mutations found in CXR clones. Nucleotidic changes and amino acid modifications are indicated for each mutated gene.

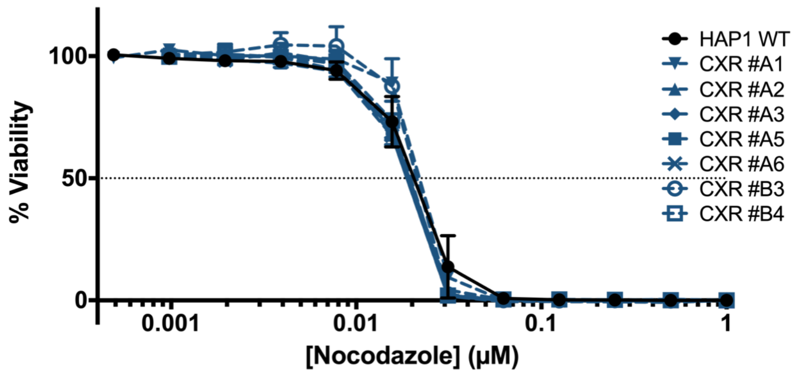


**Supplementary Figure 1a**

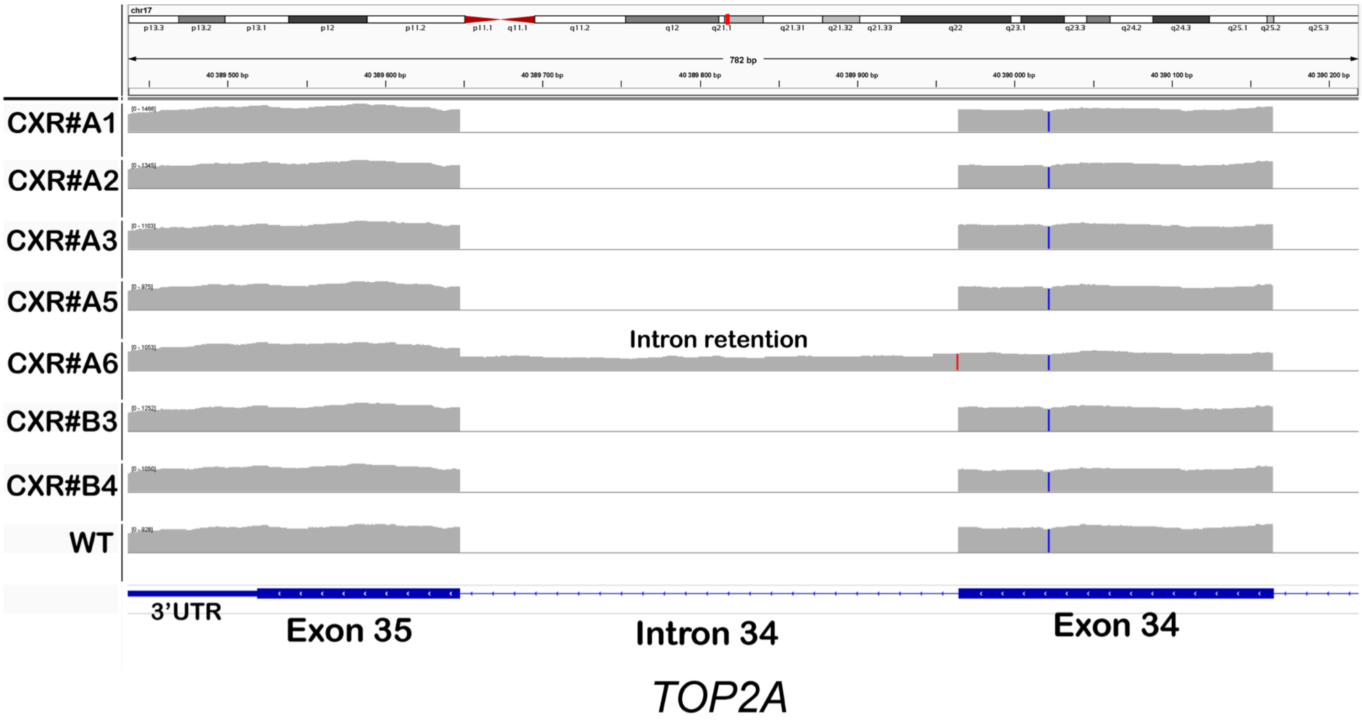


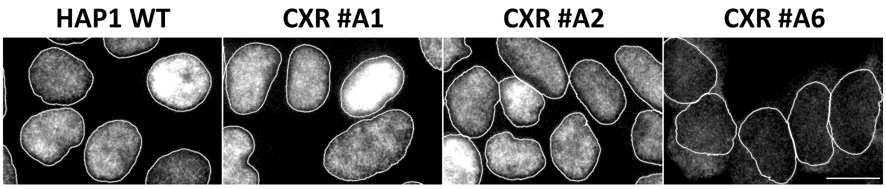


**Supplementary Figure 1b**

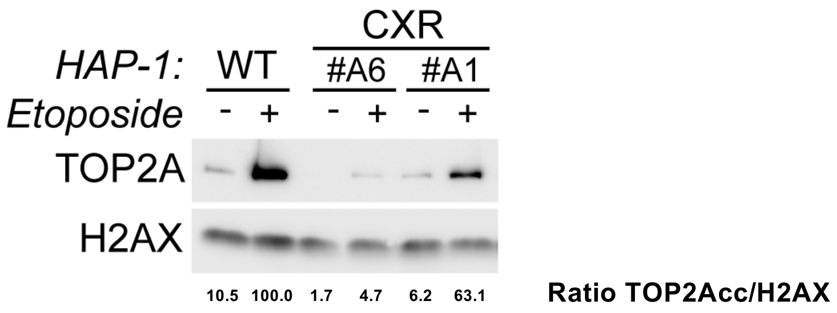


**Supplementary Figure 2**

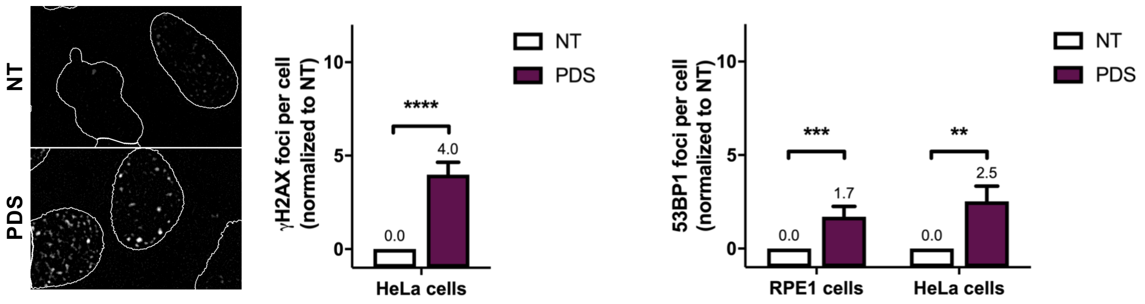


### Supplementary Figure 3a

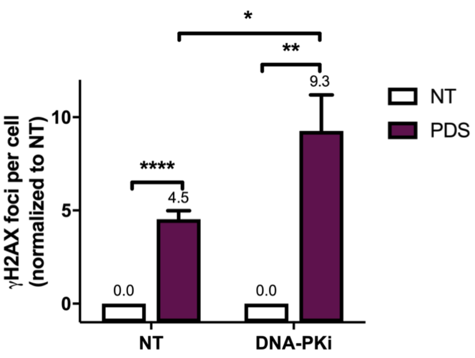


### Supplementary Figure 3b

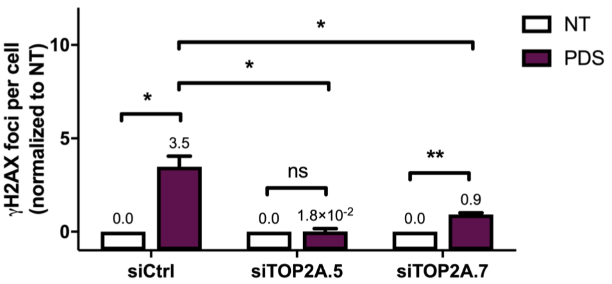


**Supplementary Figure 3c**

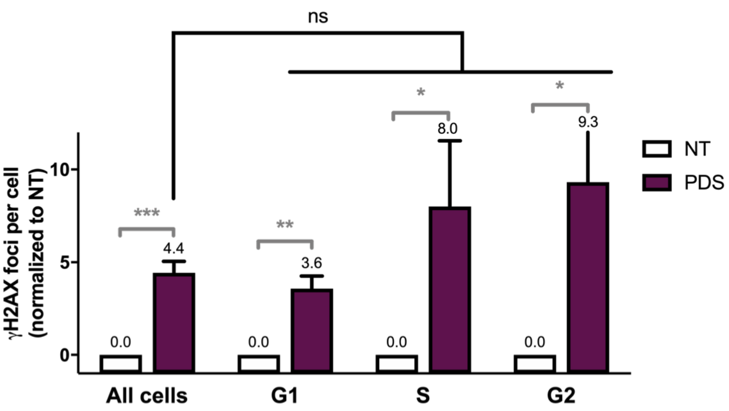


#### Supplementary Figure 4a

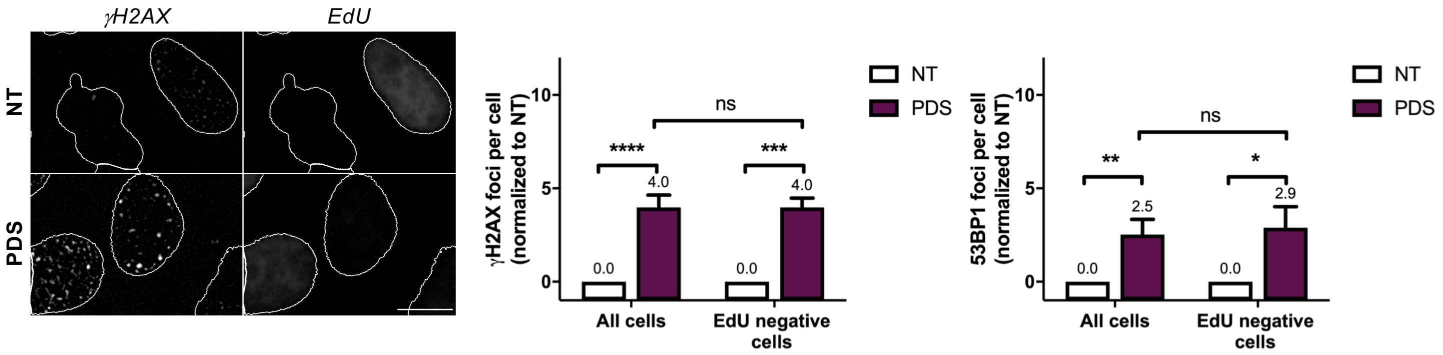


### Supplementary Figure 4b

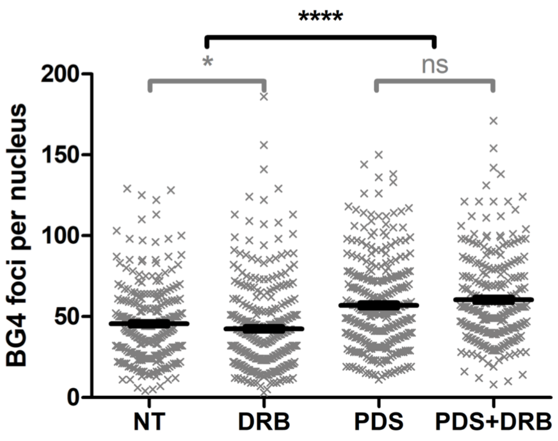


**Supplementary Figure 4c**

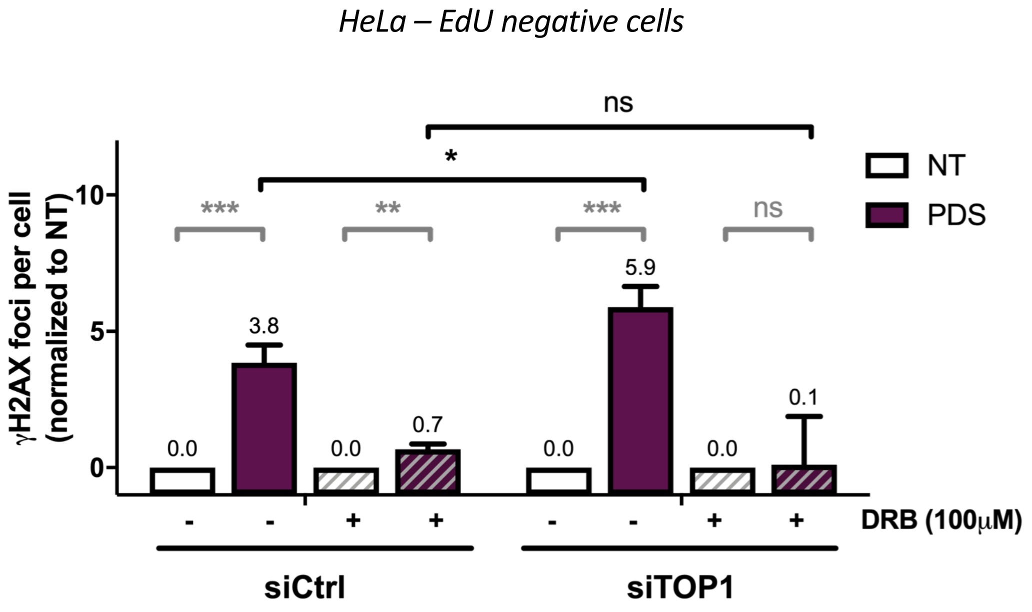


**Supplementary Figure 5**

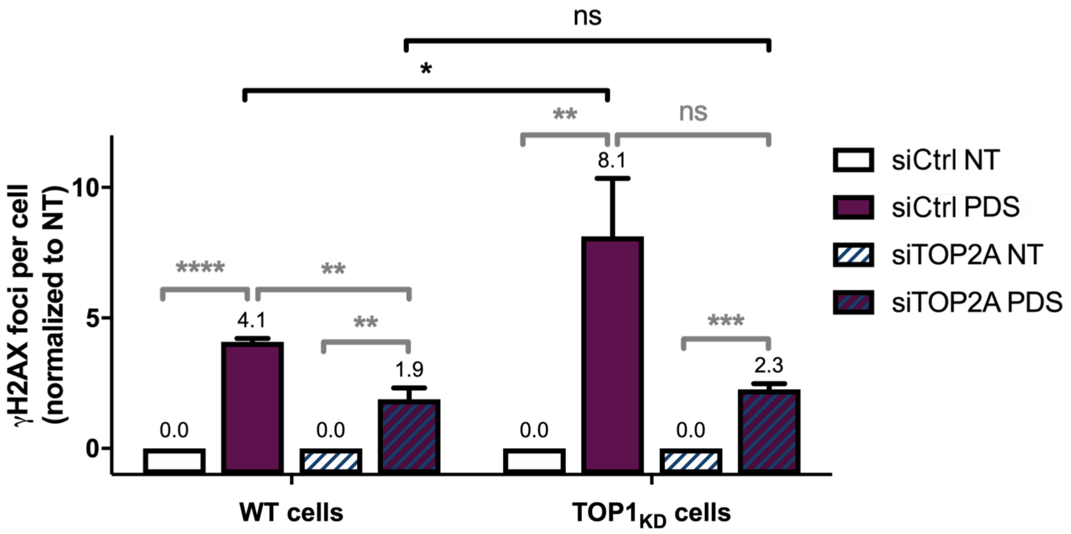


**Supplementary Figure 6**

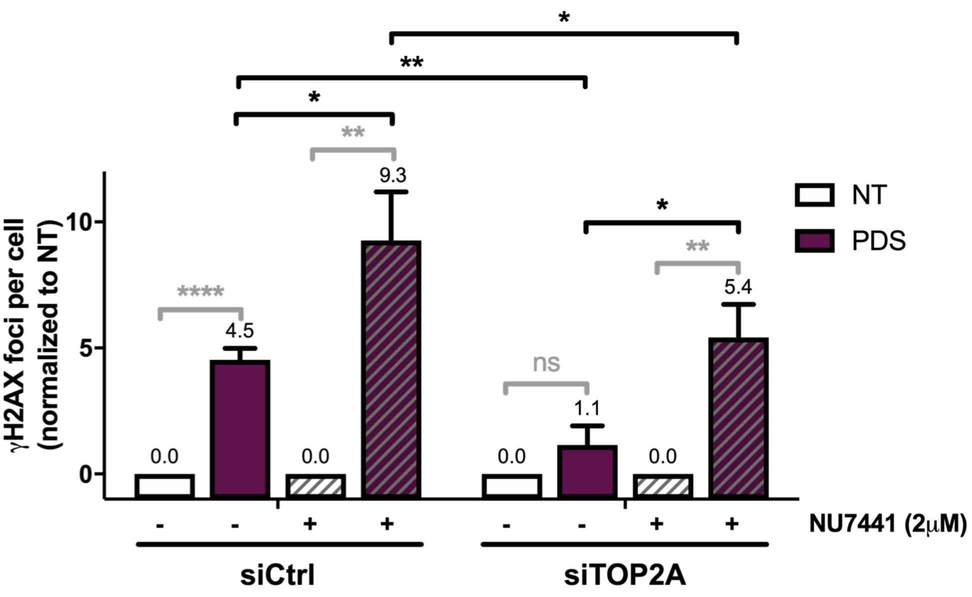


**Supplementary Figure 7**

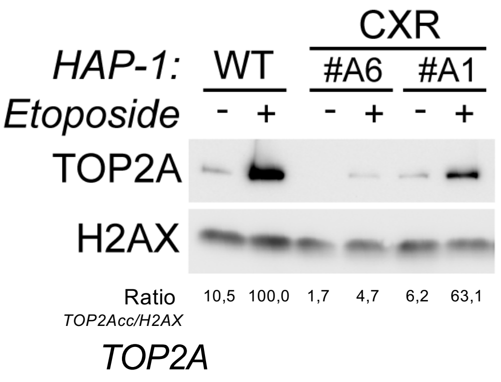


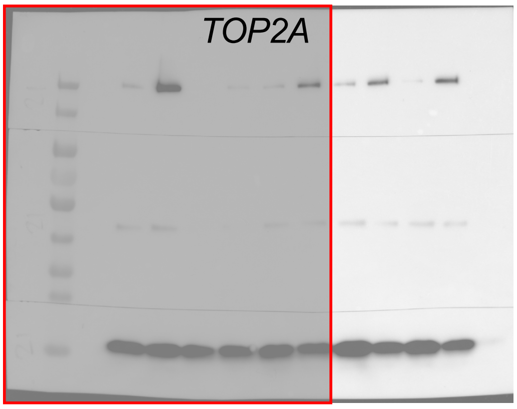


#### *Western Blot analysis from Supplementary figure 2*

Right panel represents full-length un-cropped blots of the indicate figure. Specific antibody signals are indicated for each protein. Red square represents the approximate cropped region displayed in the article

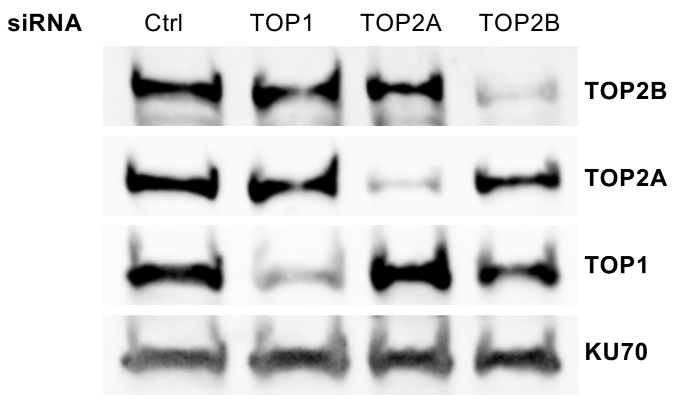


#### ***Western Blot analysis from figure 2c and 3b***

Right panel represents full-length un-cropped blots of the indicate figure. Specific antibody signals are indicated for each protein. Red square represents the approximate cropped region displayed in the article

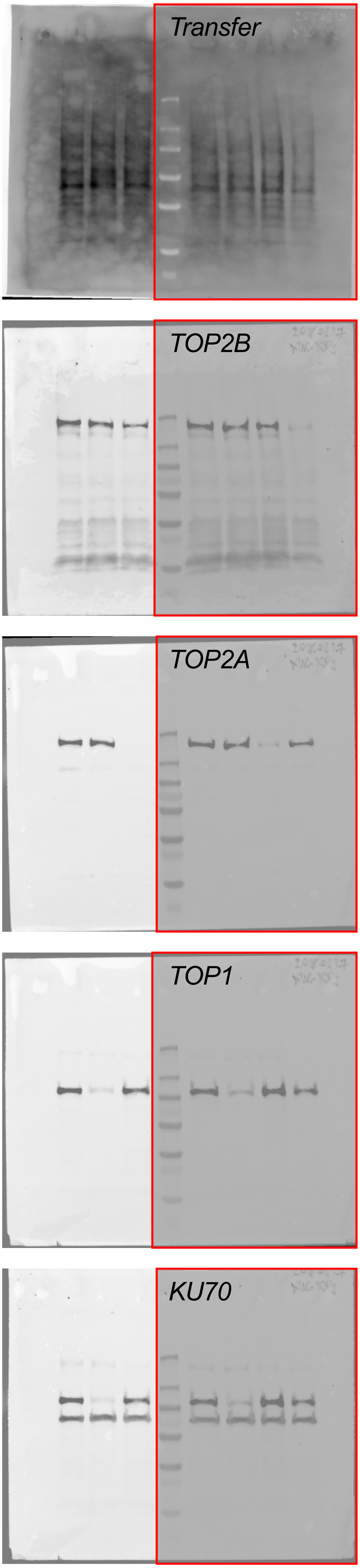

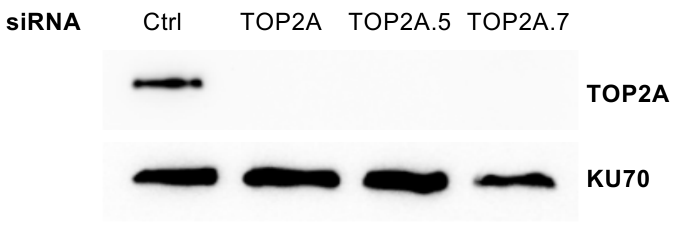


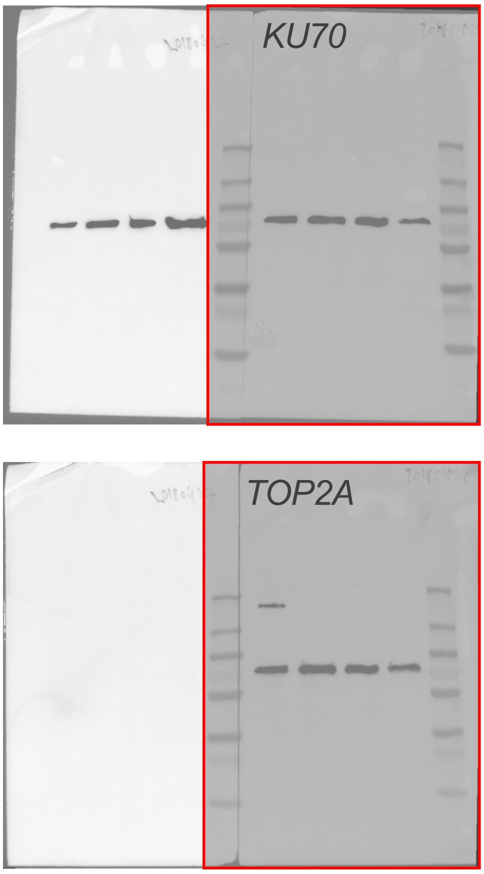


***Western Blot analysis from Supplementary figure 3c***

Right panel represents full-length un-cropped blots of the indicate figure. Specific antibody signals are indicated for each protein. Red square represents the approximate cropped region displayed in the article

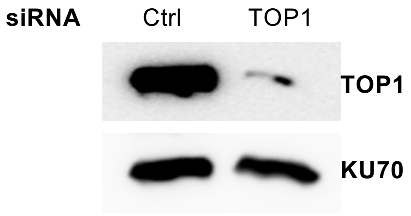


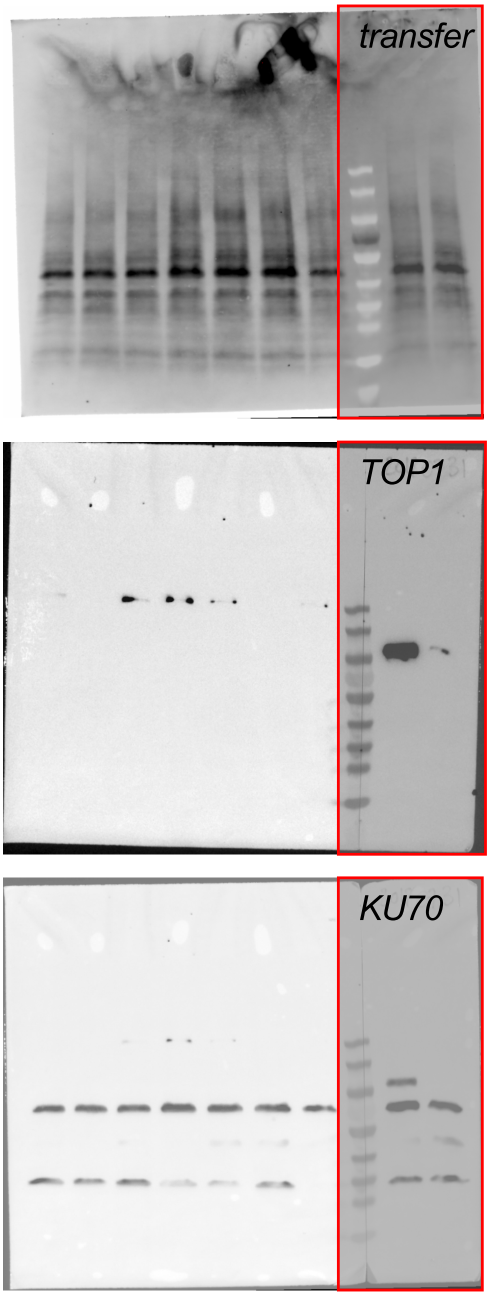


***Western Blot analysis from figure 5b, 5c and Supplementary figure 5***

Right panel represents full-length un-cropped blots of the indicate figure. Specific antibody signals are indicated for each protein. Red square represents the approximate cropped region displayed in the article

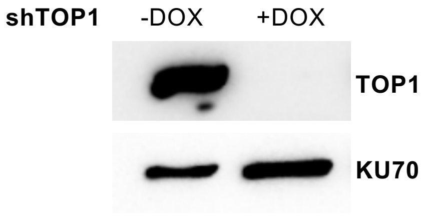


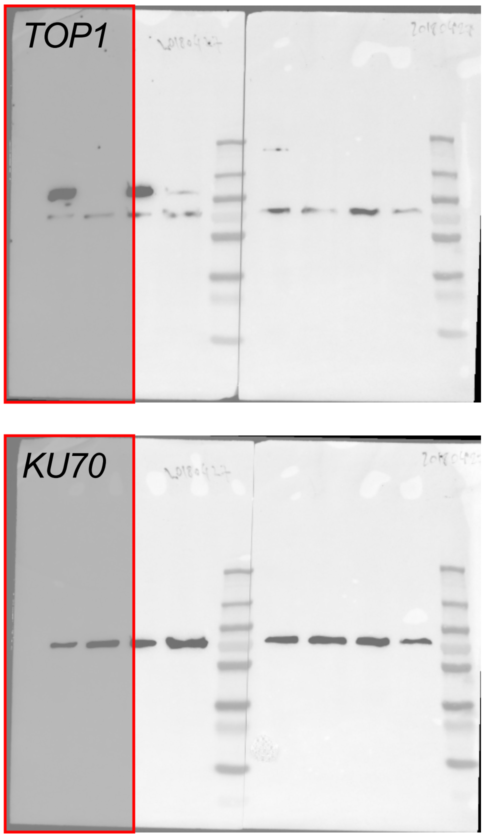


***Western Blot analysis from figure 5a, 5d and 5e***

Right panel represents full-length un-cropped blots of the indicate figure. Specific antibody signals are indicated for each protein. Red square represents the approximate cropped region displayed  in the article

**Supplementary figures legends**

**Supplementary Table 1:** $IC_{50}$ values as assessed by cell survival assays of nocodazole, etoposide, F14512 and G4 ligands (CX5461 and PDS) on WT, CX5461 resistant (CXR) and F14512 resistant (F14R) HAP1 cells.

**Supplementary Table 2:** non- or mis-sense mutations found in CXR clones. Nucleotide changes and amino acid modifications are indicated for each mutated gene

**Supplementary Figure 1a:** Viability assay of CX5461 resistant cells (CXR) to the nocodazole a microtubule binder, substrates for efflux-pumps.

**Supplementary Figure 1b:** Graphic representation of TOP2A intron retention in clone CXR#A6 carrying a homozygous mutation of the last nucleotide of the last intron, resulting in replacement of the 42 last TOP2A amino acids, carrying its nuclear localisation signal (NLS), by 18 unrelated amino acids (Top panel). Immunofluorescence analysis showing the large cytoplasmic localisation of TOP2A protein in CXR#A6 compared to WT, CXR#A1 (TOP2A F85I) and CXR#A2 (TOP2A S654I) cells (bottom panel). TOP2A staining was obtained as described in Methods.

**Supplementary Figure 2:** Heparin-based extraction protocol to monitor by immunoblotting the accumulation of TOP2Acc following ETP treatments (See Methods). In this assay, TOP2 not covalently attached to DNA is extracted by heparin in a soluble fraction, while TOP2cc is resistant to this procedure and can be analysed after centrifugation by immunoblotting of pellet fraction. As show point mutations present on clones CXR#A6 (TOP2A with a different C-terminus) and CXR#A1 (TOP2A F85I) decreased the amount of TOP2Acc. Relative protein levels of TOP2A were quantified, normalized to H2AX level, and set to 100 in WT cells treated with etoposide.

**Supplementary Figure 3a:** Quantification and representative images of γH2AX and 53BP1 foci fluorescence signal (grey) detected in HeLa cells and in RPE1 cells treated with PDS (20 μM) for 4 hours. Quantification of γH2AX foci per cell was performed as described in Methods on $n > 130$ nuclei for each condition. Quantification of 53BP1 foci per cell was performed as described in Methods on $n > 169$ nuclei for each condition in HeLa cells and on $n > 221$ nuclei for each condition in RPE1-hTERT cells. Error bars represent SD from the means, $n \geq 3$ independent experiments. *P* values were calculated using a unpaired multiple Student's *t* test. ns: $p>0.05$; *: $p<0.05$; **: $p<0.01$; ***: $p<0.001$; ****: $p<0.0001$.

**Supplementary Figure 3b:** Quantification of γH2AX in HeLa cells treated with PDS (20 μM) for 4 hours and the DNA-PKcs inhibitor (DNA PKi) NU7441. NU7441 (2 μM) was added 1 hour prior PDS addition. Quantification of γH2AX foci per cell was performed as described in Methods on $n > 147$ nuclei for each condition. Error bars represent SD from the means, $n \geq 3$ independent experiments. *P* values were calculated using a unpaired multiple Student's *t* test. ns: $p>0.05$; *: $p<0.05$; **: $p<0.01$; ***: $p<0.001$; ****: $p<0.0001$.

**Supplementary Figure 3c:** Quantification of γH2AX in HeLa cells transfected with control (Ctrl), or two different TOP2A siRNAs and treated by PDS (20 μM) for 4 hours. Quantification of γH2AX foci per cell was performed as described in the Material and Methods section on *n* >

218 nuclei for each condition. Error bars represent SD from the means, $n = 2$ independent experiments. $P$ values were calculated using a unpaired multiple Student's $t$ test. ns: p>0.05; *: p<0.05; **: p<0.01; ***: p<0.001; ****: p<0.0001.

**Supplementary Figure 4a:** Quantification of γH2AX foci throughout cell cycle in HeLa cells treated with PDS (20 μM) for 4 hours. Determination of cell cycle staging of individual cells was performed by measuring their DNA content by fluorescence microscopy (1). Quantification of γH2AX foci per cell was performed as described in Methods on $n > 130$ nuclei for each condition. Error bars represent SD from the means, $n = 3$ independent experiments. $P$ values were calculated using an unpaired multiple Student's $t$ test. ns: p>0.05; *: p<0.05; **: p<0.01; ***: p<0.001; ****: p<0.0001.

**Supplementary Figure 4b:** Quantification and representative images of γH2AX and 53BP1 foci fluorescence signal (grey) detected in HeLa cells treated with PDS (20 μM) for 4 hours in the presence of EdU (5-ethynyl-2′-deoxyuridine). EdU staining and detection is described in Methods. Quantification of γH2AX foci per cell was performed as described in Methods on $n > 130$ nuclei for each condition. Quantification of 53BP1 foci per cell was performed as described in Methods on $n > 169$ nuclei for each condition. Error bars represent SD from the means, $n \geq 3$ independent experiments. $P$ values were calculated using an unpaired multiple Student's $t$ test. ns: p>0.05; *: p<0.05; **: p<0.01; ***: p<0.001; ****: p<0.0001.

**Supplementary Figure 4c:** Quantification of BG4 foci fluorescence signal (grey) detected in HeLa cells treated with PDS (20 μM) in the presence of RNA Pol II inhibitor DRB for 4 hours. DRB (100 μM) was added 1 hour before PDS addition. Quantification of BG4 foci per cell was performed as described in Methods on $n > 70$ nuclei for each condition. Error bars represent sem from the means, $n = 3$ independent experiments. $P$ values were calculated using an unpaired Whelch's $t$ test. ns: $p>0.05$; *: $p<0.05$; **: $p<0.01$; ***: $p<0.001$; ****: $p<0.0001$.

**Supplementary Figure 5:** Quantification of γH2AX foci fluorescence signal (grey) detected in HeLa cells transfected with control (Ctrl), or TOP1 siRNAs and treated by PDS (20 μM) in the presence of RNA Pol II inhibitor DRB for 4 hours. DRB (100 μM) was added 1 hour before PDS addition. Quantification of γH2AX foci per EdU negative cell was performed as described in Methods on $n > 94$ nuclei for each condition. Error bars represent SD from the means, $n = 3$ independent experiments. $P$ values were calculated using an unpaired multiple Student's $t$ test. ns: $p>0.05$; *: $p<0.05$; **: $p<0.01$; ***: $p<0.001$; ****: $p<0.0001$.

**Supplementary Figure 6:** Quantification of γH2AX foci fluorescence signal (grey) detected in HeLa WT or HeLa TOP1 knock-down cells (inducible shRNA) transfected with control (Ctrl) or TOP2A siRNAs and treated by PDS (20 μM) for 4 hours. Expression of shTOPI in HeLa cells was induced with doxycycline (5μg/mL) 4 days prior siRNAs transfections and maintained during transfections. PDS treatment was performed 48 hours after the second round of siRNA transfection as described in Methods. Quantification of γH2AX foci per cell was performed as described in Methods on $n > 59$ nuclei for each condition. Error bars represent SD from the means, $n = 3$ independent experiments. $P$ values were calculated using an unpaired multiple Student's $t$ test. ns: $p>0.05$; *: $p<0.05$; **: $p<0.01$; ***: $p<0.001$; ****: $p<0.0001$.

**Supplementary Figure 7:** Quantification of γH2AX foci fluorescence signal (grey) detected in HeLa cells transfected with control (Ctrl) or TOP2A siRNAs and treated by PDS (20 μM) for 4 hours in the presence of DNA-PKcs inhibitor NU7441 (2 μM, DNA-PKi). NU7441 (2 μM) was added 1hour prior PDS addition. Quantification of γH2AX and BG4 foci per cell was performed on $n > 147$ nuclei for each condition. Error bars represent SD from the means, $n = 3$ independent experiments. $P$ values were calculated using an unpaired multiple Student’s $t$ test. ns: $p>0.05$; *: $p<0.05$; **: $p<0.01$; ***: $p<0.001$; ****: $p<0.0001$.
